# Neural Latching Switch Circuits for implementing Finite-State Automata

**DOI:** 10.64898/2025.12.27.696649

**Authors:** Alexis Dubreuil, Arthur Leblois, Rémi Monasson

**Affiliations:** Université de Bordeaux, CNRS, IMN, UMR 5293, Bordeaux, France; Ecole Normale Supérieure, PSL Research, CNRS UMR8023, Sorbonne Université, Paris, France

## Abstract

Cognitive processes rely on switches between mental states. Mental states are thought to be supported by cell assembly activations as introduced by Hebb. This idea has been formalized using attractor neural networks, leading to detailed mechanistic and quantitative characterizations, allowing for confrontations with neurobiology experiments. However such a mechanistic understanding is lacking for switches between mental states. Here we introduce Neural Latching Switch Circuits (NLSC) which are composed of an attractor neural network, augmented with gate neurons allowing to program transitions between network states. We explore conditions under which such circuits emerge in artificial neural networks, and provide a quantitative description by embodying NLSC into models of binary neurons. We show how NLSC can be mapped onto the fly’s head-direction system and put forward signatures of NLSC for identifying such a structure in other brain circuits. Throughout examples, we show that NLSC are suited to implement computations relevant for sensory, motor, or more abstract types of processing. To propose a meaningful characterization of NLSC in these various contexts, we interpret them as finite-state automata, identifying attractor-states and automata-states, a first step in establishing a connection between neural and symbolic computations.

## Introduction

Cognition relies on switching between different neural states encoding behaviorally relevant variables. A manifestation of this phenomenon is evident in the motor domain whereby sequences of neural states encode the succession of motor commands necessary for realizing elaborated movements (1). In the sensory domain, switching between neural states allows neural circuits to update mental representations in the face of a stream of external stimuli. For instance animals update a mental representation of their head direction upon changes in visual or vestibular sensory inputs (2; 3). Switches between neural states are also involved in more abstract forms of cognition, as evidenced in rule-based behaviors studied in laboratory conditions, such as the Wisconsin Card Sorting Test, in which subjects have to switch from one rule to another based on sensory evidence.

Numerous models in which networks switch between states have been proposed in these specific contexts: production of motor sequences (4), modeling of the head-direction system (5; 6), retrieval of episodic memories (7), rule-switching (8), or more abstract modeling of cognitive events (9; 10). They all rely on a synaptic connectivity matrix with an asymmetric, or hetero-associative, component to control transitions from one state to another, as is the case for synfire chain models (11). Other modeling studies introduced a symmetric, auto-associative, component to the connectivity matrix, allowing for instance to control for the duration spent in each state before a transition occurs (12), to flexibly control the speed at which a sequence is produced (13), or to obtain persistent neural states in the absence of switching inputs (5; 6; 8).

In this article we introduce Neural Latching Switch Circuits (**NLSC**), an elementary neural circuit that allows to program transitions between stable neural states. It is composed of an attractor neural network (14), in which auto-associative synaptic connectivity supports sustained activations of cell assemblies (15). This dictionary of cell assemblies is coupled to a set of gate neurons whose hetero-associative connectivity with the dictionary allows to program switches between attractor states upon presentation of inputs. Using training and reverse-engineering of artificial neural networks (**ANN**), we describe conditions under which NLSC emerge, and, implementing NLSC in models of binary neurons, we establish quantitative relationships between the coding and structural properties of NLSC and the computations they implement. We show how NLSC can be mapped on the head-direction system described in the fly’s brain, and discuss other predictions to identify gate neurons in other brain circuits.

Throughout examples, we show that NLSC are suited to implement computations relevant for sensory, motor, or more abstract types of processing. In order to propose a meaningful characterization of NLSC in these various contexts, we resort to computer science and interpret them as finite-state automata, identifying attractor states and automata states, a first step in establishing a connection between neural and symbolic computations.

### 1 Programming transitions between neural states using gate neurons

In order to understand how neural circuits can implement transitions between neural states, we first considered a behavioral task designed to study the neural basis of sensory-motor transformations in rodents (16). In this task, one of two sensory stimuli is presented and after a variable delay period the animal is asked to respond to a go cue by performing one of two motor actions (Fig. 1a).

**Fig. 1:**
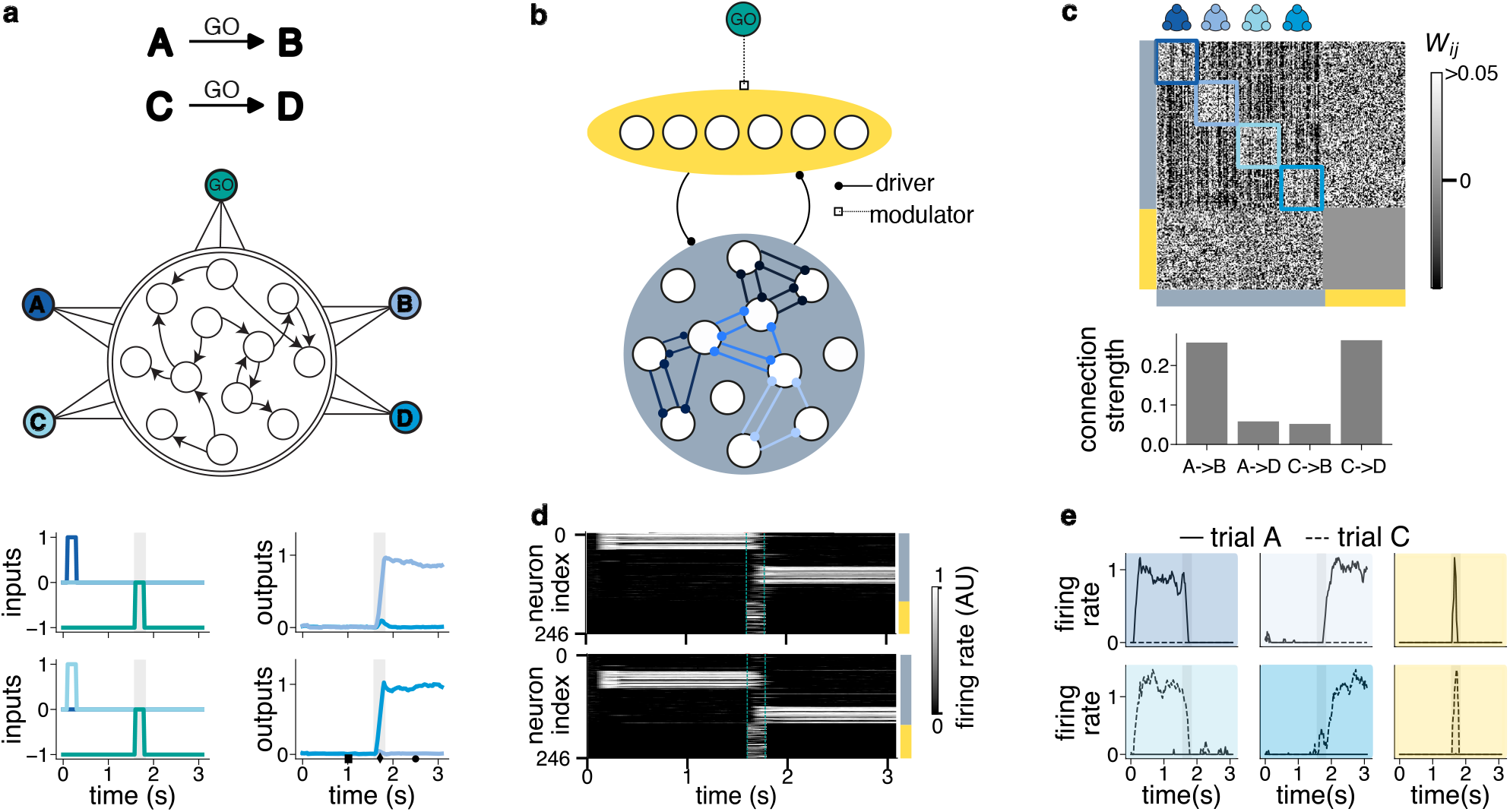
Neural Latching Switch Circuit: programming transitions between cell assembly activations with gate neurons. **a**, Top: illustration of a delayed sensory-motor transformation task, upon presentation of a go cue the system transits to a new state that depends on the previous state. Bottom: ANN trained to perform the task, with inputs and trained outputs in the two different types of trials. Grey shades represent presentation of the go input, releasing inhibition onto gate neurons. **b**, Schematic of a NLSC. Circles represent neurons. Round and squared connections represent driving and modulatory inputs respectively. A dictionary, with positive feedback loops in its connectivity implement stable cell assemblies, while gate neurons, under the modulatory control of a go cue, control transitions between cell assemblies. **c**, Top: connectivity matrix of the trained NLSC. Bottom: connectivity through gate neurons, bar size for a transition X → Y represents the averaged product of the weights from cell assembly X to gate neurons and the weights from gate to cell assembly Y. **d**, Raster of neural activity for the A to B (top) and C to D (bottom) transitions. **e**, Firing rate of six neurons for the two transitions.

We modeled results of this experiment with a neural latching switch circuit (NLSC). It is composed of an attractor neural network with four cell assemblies (Fig. 1b, blue) whose activation encodes the two sensory and the two motor states, below we refer to this part of the circuit as a dictionary of cell assemblies. Similarly to latching switch in electrical circuits, the attracting property of cell assemblies leads the circuit to latch, i.e. it remains in one of the states in the absence of inputs. Presentation of the go cue (Fig. 1b, green) allows for the activation of gate neurons through a modulatory input (Fig. 1b, yellow), leading the circuit to switch to a different state. To build NLSC, we first trained a rate network to implement the dictionary, then we added gate neurons, which are not recurrently connected with each others, and trained connections from dictionary to gate neurons and back to program task-relevant transitions (Method 7.2.1, SI Fig. 1a,c). The connectivity structure of the NLSC reflects its function, with an auto-associative component consisting of positive feedback loops in the dictionary supporting stable cell assemblies (Fig. 1c, blue squares), and an hetero-associative component through gate neurons linking sensory and motor states (Fig. 1c, bottom). Throughout task execution, individual dictionary neurons show persistent activity selective to one of the four states, while gate neurons show non-linear mixed selectivity (17), responding only to conjunction of the go cue and one of the two sensory states (Fig. 1d,e). Neural activity in the NLSC can also be described at the population level (SI Fig. 2, Methods 7.2.3,7.4), with inputs from the go cue modulating the gain of gate neurons and controlling interactions between cell assemblies (18).

### 2 Hetero-associative connections through gate neurons is more efficient than direct connections between cell assemblies

Recordings of neural activity while rats perform this task showed that sensory and motor states are encoded in pre-motor cortex, and that thalamic neurons respond to presentations of the go cue (16). If this cortico-thalamic network implements a NLSC, the current pre-motor cortical state could be decoded in the associated motor thalamus, with individual neurons exhibiting non-linear mixed selectivity as in Fig. 1e-right. We note that for this task it does not have to be the case, and thalamic circuits could also simply relay the go cue. Indeed, by analyzing 50 ANN trained without a priori architectural constraints (Method 7.2.2), we observed that transitions from sensory to motor states can in principle be programmed by directly shaping the connectivity in-between cell assemblies of the dictionary, without resorting to gate neurons (Method 7.3.2, SI Fig. 3 d-g). In a given trial, a cell assembly encoding a motor variable is correctly activated, e.g. C, if it receives more current than the other motor variable, e.g. D. This *switching* current has two sources, as represented by the dotted and dashed connections in the cartoon of Fig. 2a: direct currents coming from the preceding sensory cell assembly activation, and indirect currents coming from gate-like neurons transiently active at the go cue. For the ANNs trained without architectural contraints, we found that 83% of the difference in currents received by the two motor cell assemblies are due to direct connections from sensory cell assemblies, and that 17% of the difference in received currents originates from gate-like neurons (Fig. 2a, hatched boxes, Method 7.3.2). While these two network mechanisms are known to be able to implement transitions between neural patterns, modulating effective interactions between dictionary neurons through gate neurons, as in the NLSC (SI Fig. 2), has been shown to be more robust for triggering many transitions (19).

**Fig. 2:**
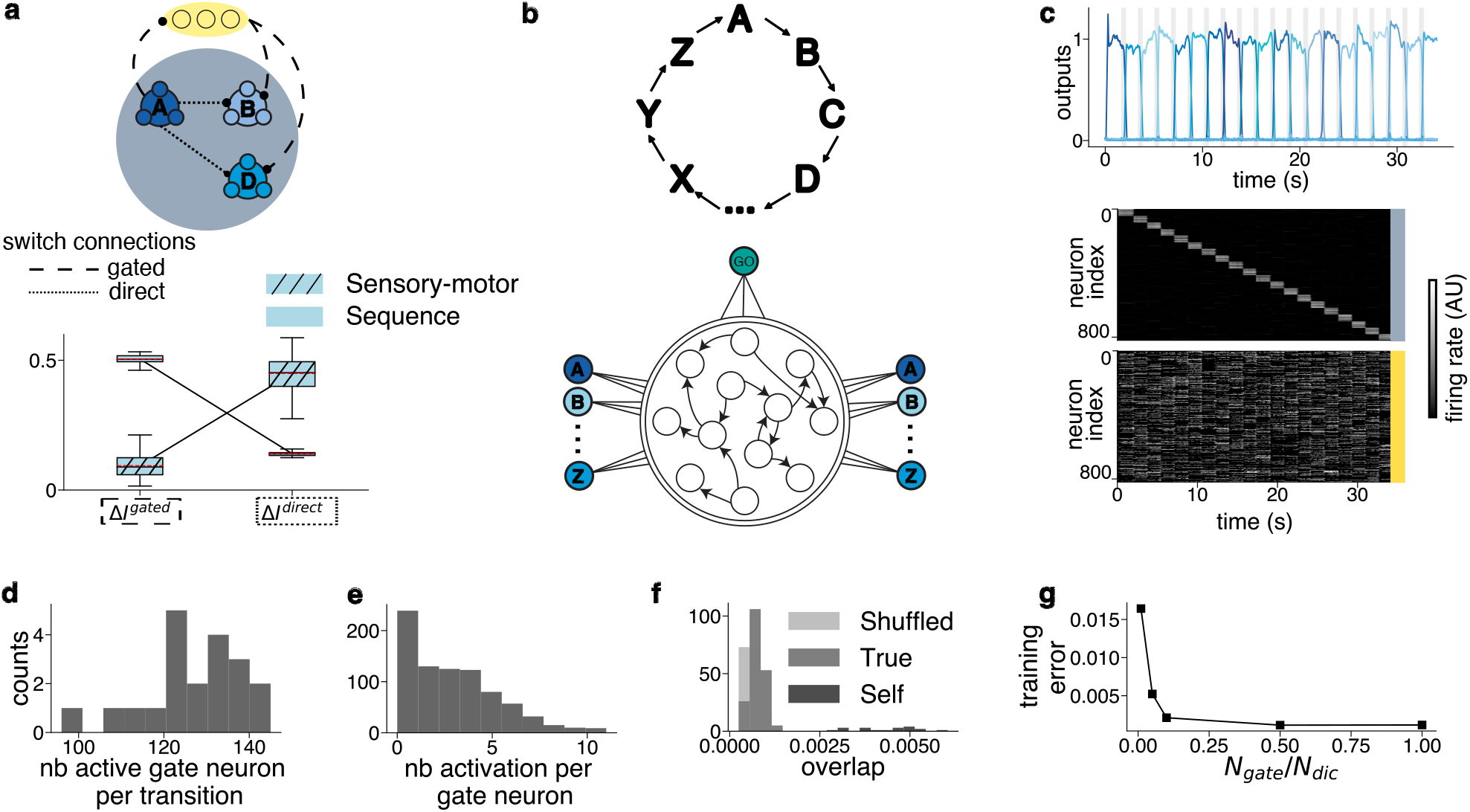
NLSC for sequence generation with gated-hetero-associative connectivity. **a**, The heteroassociative connectivity component that programs transitions between neural state can be carried by direct connections between subsequent cell assemblies (dotted lines), or through gate neurons (dashed lines), as illustrated for the A to B transition of the sensory-motor transformation task. Box plots show the proportion of direct and gated currents responsible for state switching, averaged over the two types of transitions for 50 trained ANN without a priori architectural constraints, for the sensory-motor transformation and the sequence tasks. **b**, Permutation of dictionary states and sequence generation. **c**, Sequential activity of an ANN trained to implement a permutation of 20 states. Top: activity of the readout neurons. Bottom: raster of individual neurons in dictionary (light blue) and gate (yellow) populations. **d**, Number of gate neurons active in one transition **e**, number of transitions in which a gate neuron is activated. **f**, Comparing activation patterns of gate neurons across all transitions: distributions of overlaps between all pairs of patterns. The true distribution is compared to the distribution of the self-overlap of the 20 patterns, as well as to overlaps obtained by randomly shuffling neuron identity. **g**, Training error as a function of the ratio between the numbers of gate and dictionary neurons.

Consistently, when training unconstrained ANNs to implement a permutation of 20 motor states (Fig.2b, Method 7.3.3), the difference in received currents onto the next cell assembly to be activated dropped at 22% for direct currents and increased to 78% for indirect currents (Fig. 2a and SI Fig. 4 f,g). In Fig. 2c we show the activity of a trained NLSC producing a sequence of 20 neural states, that can be thought of as a sequence of motor commands. A main goal of this study is to characterize properties of gate neurons, we thus focused on neural activation patterns of gate neurons in response to the go cue. Gate activation patterns were distributed across multiple neurons: each transition involved the activation of 129 out of the 820 gate neurons and each gate neuron was involved in 3 transitions on average (Fig. 2d,e, SI Fig. 4b). In addition gate patterns were orthogonal to each others (Fig. 2f, SI Fig. 4c), as expected from the fact that they correspond to the imprinting of orthogonal dictionary states and that they should each point towards one of these orthogonal dictionary states. We noticed that in order to implement the 20 transitions, the number of gate neurons could be reduced from 820 to 82 without loss in performance (Fig. 2g). This lead us to wonder how many neurons are required to program transitions between dictionary states. In principle, having only 20 gate neurons, one per transition is feasible (19). But given that patterns of activation of gate neurons are distributed, it could be possible to lower this number.

In order to estimate this number and to go beyond the qualitative descriptions obtained by analysis of trained ANN, we embodied NLSC in models of binary neurons and adapted previously developed mathematical techniques for studying the storage capacity of attractor neural networks or perceptrons with binary synaptic weights (20; 21; 22) (Method 7.5, Supplementary 8.1). In the spirit of (23; 24) models are optimized, for a given task, by finding the parameters that minimize the required number of neurons, *N*_*min*_. To do so we treated the different types of connections of a NLSC as three perceptron problems (Fig. 3a, SI Section 8.1). For a NLSC implementing a permutation of *P* dictionary states, *N* _*min*_ scales as 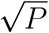 (Fig. 3b), as already known for attractor neural networks with sparse neural patterns (25; 22). A total of *N*_*min*_ ≃ 10, 000 neurons, the approximate size of a cortical column, allows a NLSC to implement a permutation of *P* ≃ 33, 000 dictionary states, with dictionary and gate patterns characterized by a coding level *f* ≃ 0.003, i.e. 0.3% of dictionary or gate neurons are activated in each pattern of activity or at each transition. The fraction of the binary synapses that are potentiated, inside the dictionary or in between dictionary and gates, is 0.15 (SI Fig. 7e).

**Fig. 3:**
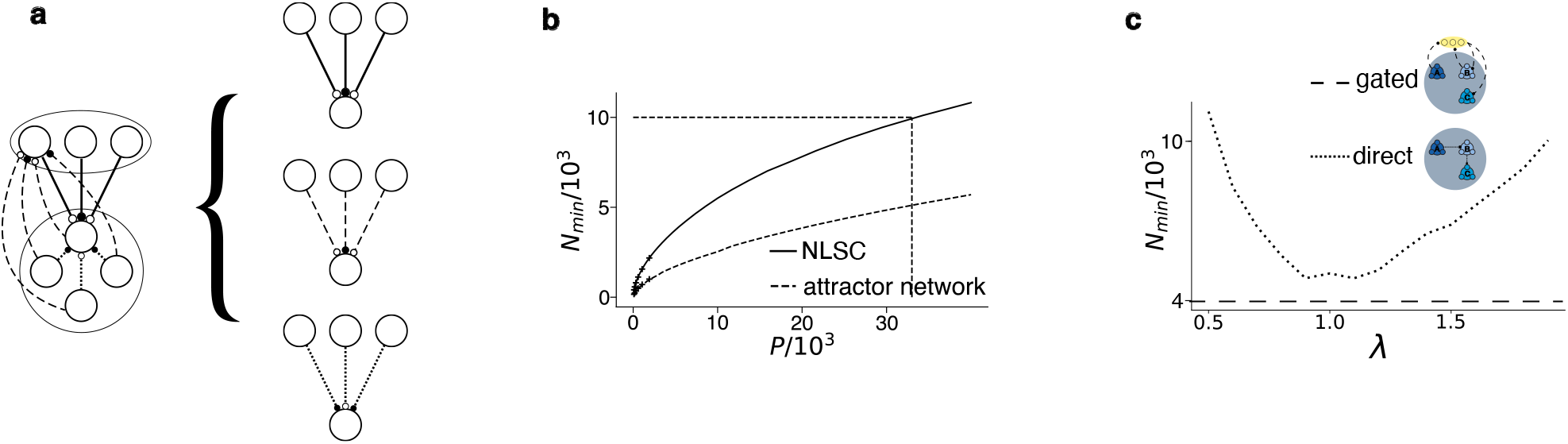
Minimal number of neurons to implement a permutation of dictionary states. **a**, The calculation of N_min_ relies on treating the dictionary and gate neurons as inputs and/or outputs of perceptrons. **b**, N_min_ for an attractor neural network storing P patterns, or a NLSC implementing a permutation of P dictionary states. Lines are obtained from analytical calculations, while crosses represent simulations results. **c**, N_min_ for a NLSC and a model with direct-hetero-associative connectivity implementing a permutation of P = 5, 000 dictionary states.

By contrast, the network mechanism without gate neurons is less efficient. This is shown in Fig. 3c, where we compare a NLSC with a direct model, in which the same synapses implement both the auto-associative component for stable cell assembly activations and the hetero-associative component for programming switches between them (cartoon Fig. 2b, SI Sections 8.1.3-8.1.4). The parameter *λ* weights the importance of these two types of structure on the synaptic connectivity for the direct model. We found that for all values of this parameter, *N*_*min*_ is smaller for the NLSC model. Thus, using gate neurons minimizes overall interferences by representing cell assembly stability and cell assembly switches onto different sets of synapses. This results is consistent with numerical estimates which show that for producing long sequences, using a single gate neuron per transition leads to more robust networks than those using direct connections between cell assemblies (19).

### 3 NLSC as finite-state automata

The sensory-motor transformation task (Fig.1a), or the production of sequences of motor states (Fig.2b), can be modeled by finite-state automata (**FSA**, Fig.4a). FSA describe machines programmed to carry out simple tasks, such as running an elevator. The computations they perform are depicted as a succession of machine states, with sensory inputs from the environment triggering state transitions, and some of the states acting on the environment through activation of effectors (26). The behavioral tasks we have thus far implemented with a NLSC are modeled by a specific type of finite-state automata, sequential machines, which are characterized by a single sensory input playing the role of a clock, as in an electrical circuit running a traffic light.

To go beyond behaviors modeled by these simple FSA, we now focus on the task of head-direction encoding, modeled by the FSA in Fig. 4b. Such a FSA is called a categorizer: all sequences of the two clockwise/counter-clockwise vestibular inputs are categorized into an angular position represented by one of the state of the automaton. We trained ANN to maintain a representation of their current angular position, as well as to update this position in response to activation of one of the two vestibular velocity inputs (Fig. 5a, Method 7.3.4). We first trained unconstrained ANN on this task and confirmed that networks resorted to gate neurons to trigger transitions between cell assemblies encoding head-directions (SI Fig. 5f,g). Gate neurons were not necessary to implement sequential machines with a single external input, although they lead to a better efficiency of NLSC (Fig. 3c). Here instead, we argue that gate neurons are necessary, to reformat the neural representations of the dictionary and the two velocity neurons into linearly separable ones. In Fig. 5c we show how the task of head-direction encoding boils down to a XOR problem. In Fig. 5d we illustrate how gate neurons can play the role of a hidden layer mixing dictionary and velocity neural representations. This is a similar role to the one proposed for gate neurons in a model of rule switching (8).

**Fig. 4:**
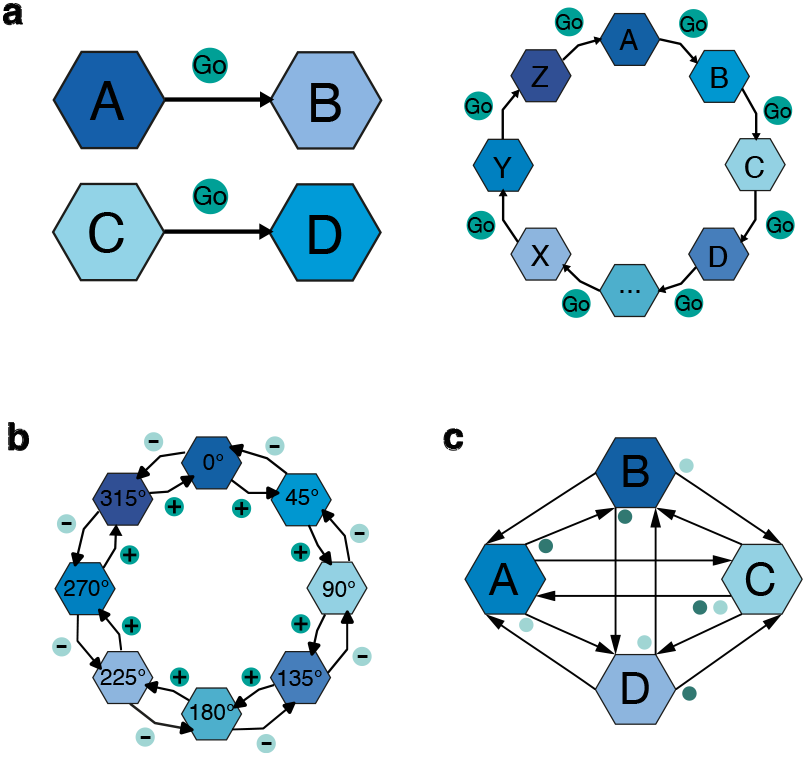
Examples of finite-state automata. Blue hexagons represent machine states, arrows represent states transitions in response to external inputs represented as green dots. **a**, Sequential machines for solving the sensory-motor transformation task (left) and generating sequences of 20 states (right). **b**, Categorizer implementing a head-direction system. **c**, FSA describing a machine with 24 external inputs, each driving a permutation of 4 states, not all inputs are shown.

**Fig. 5:**
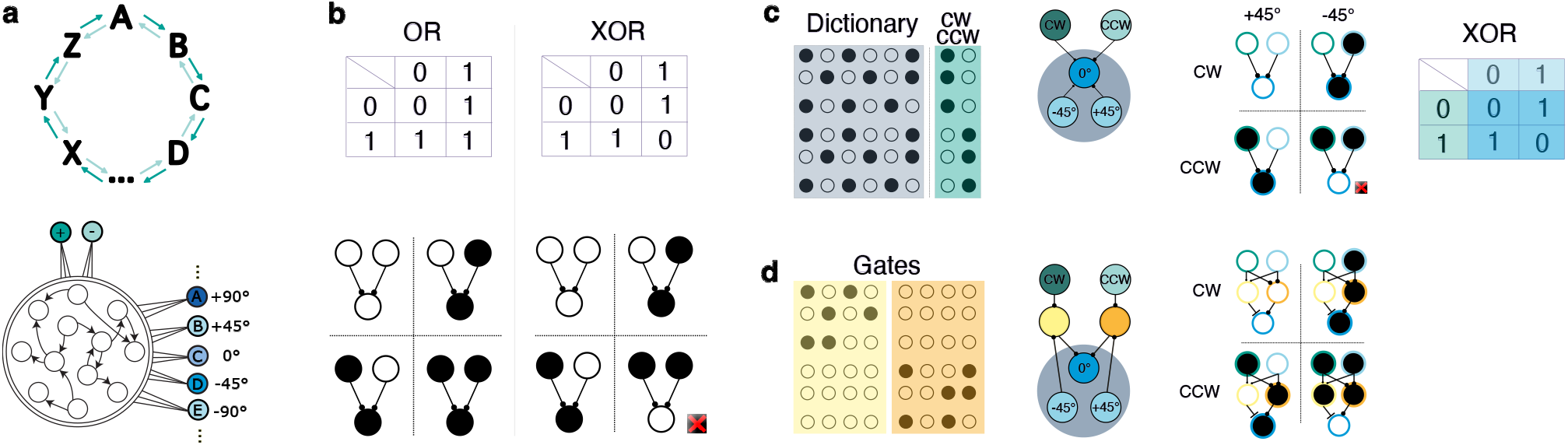
Head-direction task as a XOR problem. **a**, Schematic of a neural network, with internal states representing angular positions, under the control of clockwise and counter-clockwise vestibular velocity inputs.**b**, Truth tables for the OR and XOR problems (top), illustrations with simplified neural representations (bottom), active neurons are black and silent neurons are white. A perceptron with an output neuron with an activation threshold of 0.5, receiving from two weights of 1, solves the OR problem. A perceptron cannot solve the XOR problem. **c**, The head-direction task requires to solve a XOR problem. Left: Each line represents an input neural pattern that has to be associated with future activation of dictionary neurons. Blue shaded area represents the part of the input neural pattern carried by the dictionary neurons, green shaded area represents the part of the pattern carried by the velocity inputs neurons. Right: simplified neural representations of the input neural patterns. Activation of the left input neuron (green circled) indicates which of the two velocity neurons is active, activation of the right input neuron (light blue circled) indicates which of the +45° or -45° cell assembly is currently activated in the dictionary, activation of the output neuron (dark blue circled) indicates whether or not the 0° cell assembly should be activated next. **d**, Left: linearly separable gate activation patterns. Right: simplified neural representations as in **c**, with the addition of the yellow and orange hidden neurons.

### 4 Gate neurons are organized into populations

We then trained a NLSC on this task and examined the emerging structure of activity in gate neurons. Examination of the patterns of activations (Fig. 6b bottom, green delimiters) for all transitions revealed that gate neurons segregated into two populations, one population of neurons active during clockwise turns and another one active during counter-clockwise turns (Fig. 6c, SI Fig. 5c,d,e). This maps well onto the two populations of the protocerebral bridge in the central complex of the fly’s brain, with neurons exhibiting non-linear mixed selectivity, mixing angular position and vestibular velocity in one direction (3). The emergence of these two populations of gate neurons, each responding to a single vestibular cue, can be understood as a particular way to have linearly separable gate patterns to point to the correct next dictionary state (Fig. 5d).

**Fig. 6:**
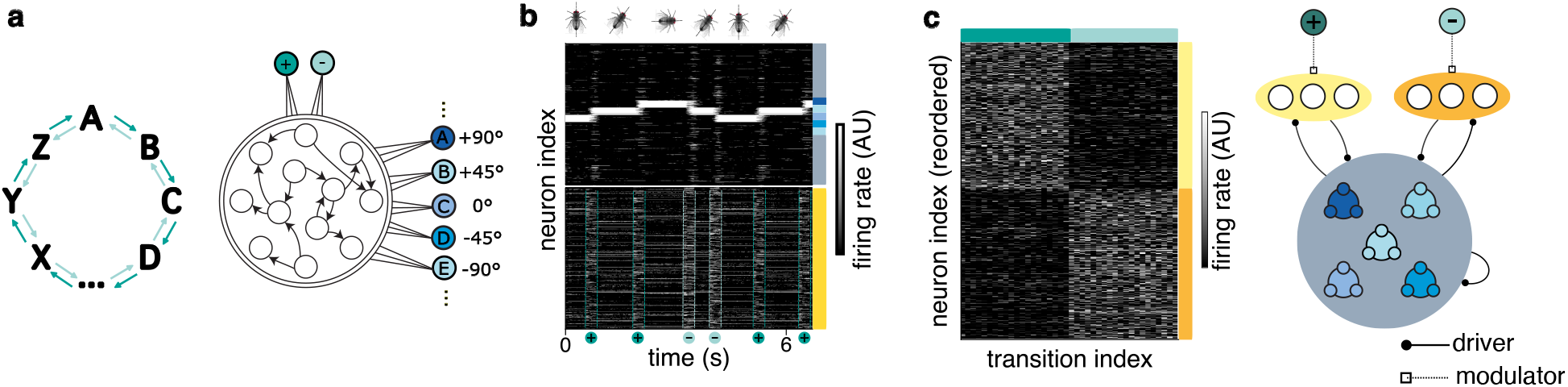
NLSC implements a head-direction system. **a**, Modeling head-direction encoding as two inverse permutations of dictionary states. **b**, Raster of neural activity, in a trained NLSC with 20 dictionary states, for a sequence of turns (green dots). Top: activation of cell assemblies in dictionary neurons, bottom: activations of gate neurons. **c**, Left: Raster of patterns of activation of the 820 gate neurons, for the 20 clockwise (left part of the X-axis) and 20 counter-clockwise transitions (right part of the X-axis). Right: Schematic of the NLSC for the head-direction task.

In order to better understand how NLSC can implement FSA, we explored another case with multiple external inputs (Fig. 4c, Fig. 7a,b), but this time their number *P*_*c*_ was much higher than the number *P* of dictionary states (while *P*_*c*_ = 2 and *P* = 20 in the head-direction example). With dictionary states corresponding to motor commands, this would model the production of all possible motor sequences with a set of effectors. Training a NLSC to implement all *P*_*c*_ = 24 permutations of a dictionary with *P* = 4 states required to correctly scale input connectivity weights with respect to dictionary-gates weights, as well as to increase the number of gate neurons. We found this time that gate neurons segregated into 4 populations (Fig. 7c-left, SI Fig. 5j-l, Method 7.3.4). In the head-direction case, external inputs were modulating gate neurons by controlling their gain (maintaining them below firing threshold) and dictionary neurons were driving them (27) (Fig. 6e-right). Here we observe the opposite with dictionary states controlling the gain of gate neurons and external inputs driving them (Fig. 7c-right). This other form of mixing is another way to obtain linearly separable gate patterns.

**Fig. 7:**
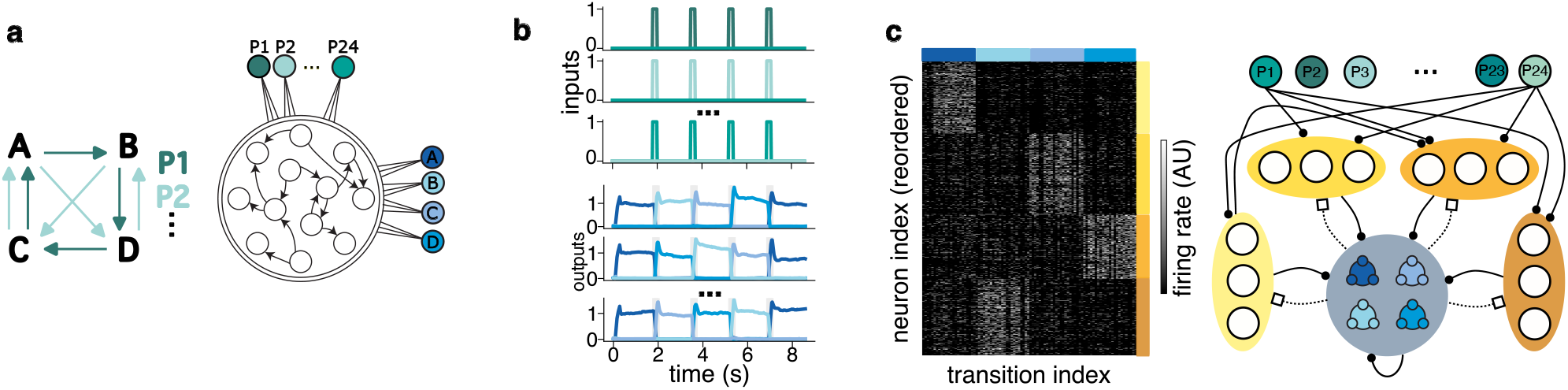
Implementing all permutations of a small set of dictionary states. **a**, Training networks to implement 24 permutations of a set of 4 dictionary states. **b**, Activation of readout neurons (bottom, blue) upon transient activations of one of the external inputs (top, green) (3 example permutations). **c**, Left: Patterns of activation of the 1640 gate neurons. Transitions are ordered on the X-axis to first display the 24 transitions starting from state A, then those starting from states B, C and D. Right: Schematic of the NLSC for all permutations.

### 5 Computational capacity of NLSC

To understand when it might be better to structure gate neurons of the NLSC as in Fig. 6c or Fig. 7c, we compared values of *N*_*min*_ in models of binary neurons for these two implementations, under different values of *P* and *P*_*c*_. Consistent with the organization emerging in ANN, we found that when *P*_*c*_ ≪ *P*, splitting gate neurons into *P*_*c*_ populations lead to smaller values of *N*_*min*_ compared to splitting into *P* populations. The converse was true for *P*_*c*_ ≫ *P* (Fig. 8a, dashed lines). This can be understood by considering the need to demix the two sources of inputs onto gate neurons and the need to use sparse-coded neural patterns to minimize ressource use (28) (SI Section 8.1.5): it is more economical to have order 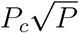 gate neurons rather than order 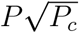 in the case *P*_*c*_ ≪ *P*, and the converse for *P*_*c*_ ≫ *P*.

**Fig. 8:**
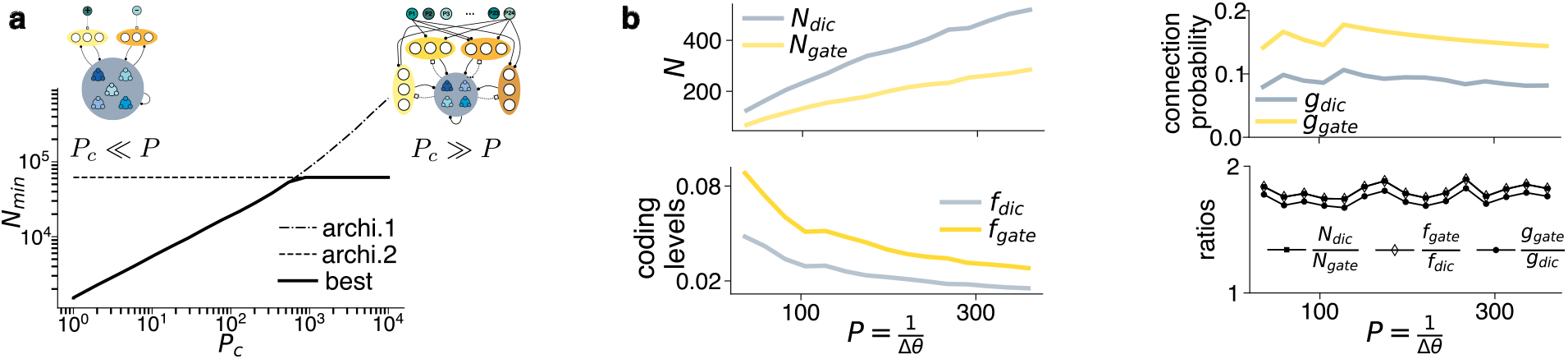
Computational capacity of NLSC. **a**, Computational capacity of NLSC implementing FSA. In the first architecture (dotted-dashed line), external inputs define P_c_ populations onto gate neurons through gain modulation, while in the second architecture (dashed line), dictionary states define P populations onto gate neurons. **b**, Structural and coding characteristics of NLSC optimized to implement head-direction systems with improving resolutions.

Studies on attractor neural networks defined the notion of capacity of a neural architecture, which measures how the minimal amount of ressources (e.g. number of neurons) depends on computational demand (e.g. number of stable cell assemblies) (29). We characterized the computational capacity of the NLSC by the minimal number *N*_*min*_ of neurons required to implement a FSA specified by its number of machine states *P*, number of possible input symbols *P*_*c*_ and fraction *E* of existing transitions from one machine state among the *P*_*c*_ possible ones (SI Section 8.1.3). The full line in Fig. 8a shows *N*_*min*_ as a function of the number of input symbols *P*_*c*_ for *P* = 1000 machine states and *ϵ* = 1. Our analytical approach allows to characterize optimal neural architectures by their number of dictionary and gate neurons, number of gate populations, organisation in terms of driver and modulator inputs, coding levels of activity patterns in dictionary and gate neurons, and average fraction of active synapses between structures (SI Fig. 7i,j).

We illustrate this by considering NLSC optimized to implement head-direction systems. Head-direction systems have been described in various animal models (30; 2; 31), with different angular resolutions, as suggested by the observation of wider bumps of activity in zebrafish larvae compared to flies. For decreasing angular resolutions 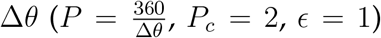, *N*_*min*_ goes from 193 to 805 for Δ*θ* = 10° to Δ*θ* = 1°. The coding levels of dictionary and gate patterns decrease with decreasing resolution, the average connectivities remain constant, and a ratio of 1.8 between quantities for dictionary and gates is kept constant (Fig. 8b). It would be interesting to see whether the head-direction system of the zebrafish larvae (31) also relies on gate neurons, and to compare their properties with those of flies (2). We note that even for a low Δ*θ* = 10° resolution, *N*_*min*_ = 193 is reached for a large number of neurons compared to the number observed for the fly’s brain. In (3), the authors estimated that 54 neurons support attractor states and that each population of gate neurons in composed only of 9 neurons. An assumption in our modeling set-up is that the fixed-points we considered in dictionaries are randomly dispersed throughout state space, as opposed to models of continuous attractors, such as the ring model which is typically used to model the head-direction system. As pointed at in the discussion, it would be interesting to extend our quantitative analysis to the case of continuous attractor networks and to describe bump motion (32) in small head-direction models (3; 33).

### 6 Oscillatory activity driving state update

In the examples we have considered so far, transitions between cell assembly activations in NLSC are triggered by an external go cue. We wondered how such transitions could be triggered internally. We trained a NLSC to autonomously switch between its dictionary states every one second (Fig. 9a). To do so, the network developed an oscillatory mode (Fig. 9b,c), with patterns of gate activations triggering transitions among dictionary states (SI Fig.6a,b). At the network level, dictionary states became meta-stables (34). While at the single cell level, this oscillatory mode manifested as dictionary neurons having ramping up or ramping down activations (Fig. 9d,e,f), as observed in singing birds (35) or in working memory tasks with delays with a characteristic time scale (36; 37; 38).

**Fig. 9:**
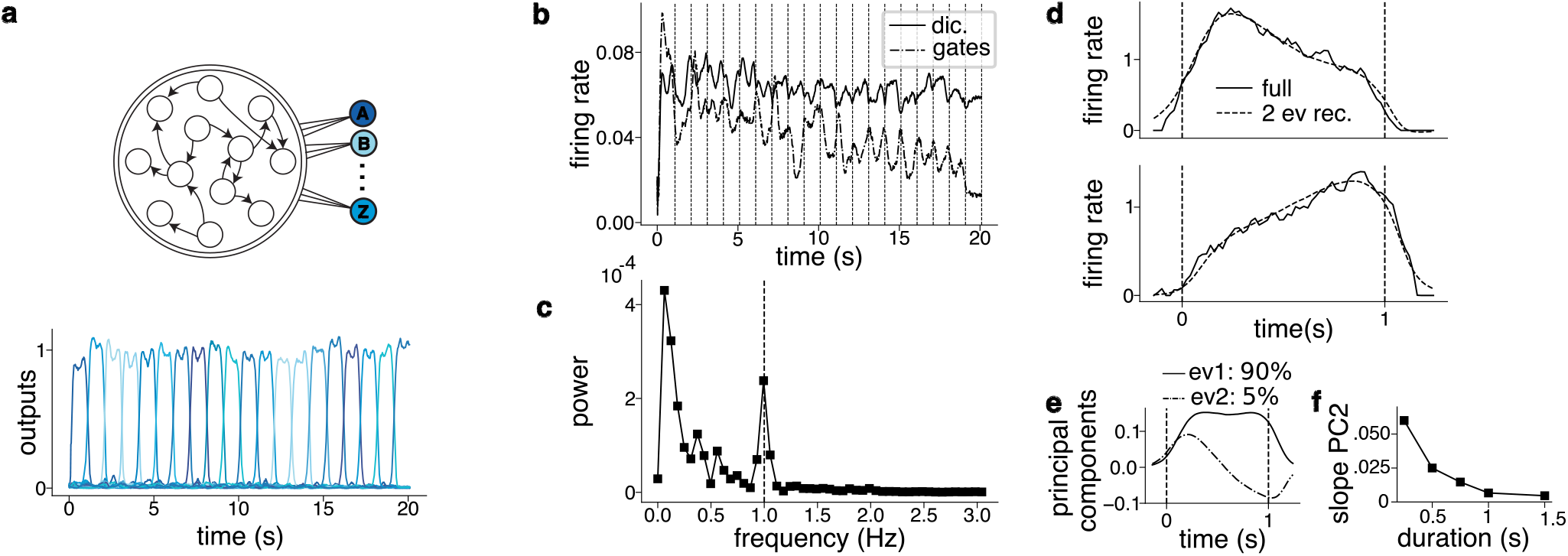
Oscillatory activity driving state update. **a**, ANN trained to autonomously switch states every one second. **b**, Average firing rate of dictionary and gate neurons. **c**, Power spectrum of the average activity of gate neurons. **d**, Firing rate of two example dictionary neurons. **e**, Temporal profiles of dictionary neurons are well explained by two components. **f**, Slope of the second ramping component as a function of state duration.

## Discussion

In this paper, we have introduced Neural Latching Switch Circuits (NLSC) which are neural network implementations of Finite-State Automata (FSA) (26). They are composed of an attractor neural network (14) which enables the activation and maintenance of multiple cell assemblies (15) encoding machine states. The attractor network is augmented with gate neurons that allow to program transitions between machine states upon presentations of inputs. Experimenting with artificial neural networks trained on simple tasks, we have outlined conditions under which NLSC emerge, and characterized the emerging population structure in gate neurons. We have implemented NLSC in models of binary neurons, and extended previous capacity calculations characterizing the ability of perceptrons to store input-output associations or of attractor neural networks to maintain persistent neural patterns (see e.g. (29; 20; 21; 22; 39; 40)). This has allowed us to define the computational capacity of NLSC and to quantitatively describe how the coding and structural properties of NLSC can be optimized to meet specific computational demands.

### Using gate neurons is efficient to implement sequences of neural states

We have studied the production of short and long sequences in ANN. We observed that while for short sequences the hetero-associative component was carried mainly by direct connections from one cell assembly to another, for longer sequences it was carried mainly by connections through gate-like neurons. Those observations are consistent with simulations from (19) which compare sequence production with both types of implementation and show that gated hetero-associative connectivity are better suited for longer sequences. We embodied both mechanisms in models of binary neurons and showed that NLSC need less neurons than a model with direct-hetero-associative connectivity to store long sequences of neural patterns. In the study of (19), each gate neuron is associated to a single transition. Instead, in trained NLSC we observed that transitions are associated with distributed patterns of gate activations and that a given gate neuron is involved in multiple transitions. Studying the binary model implementation of NLSC lead us to show that such distributed representations, which can leverage the advantage of sparse coding (41; 42; 28), are beneficial as they require a number of gate neurons that scales as the square root of the number of independent transitions, not as the number of transitions.

Other models of sequence production have proposed to use gate-like neurons to segregate auto- and hetero-associative components onto distinct sets of synapses. For instance, these models allowed to account for the duration distributions of behavioral states in a self-paced behavior (43), or to propose a mechanism to control the production speed of a sequence of neural patterns (13), although in this case a clear segregation between gate and dictionary neurons was not necessary. Another study, modeling the generation of sequences of motor commands (44), proposed to use dictionary and gate neurons, allowing a network to re-use dictionary patterns in the production of different neural sequences.

### Population structure of gate neurons depends on computational requirements

Such a re-use of neural states in different sequences is also required for the processing of sensory input streams. A canonical example is the head-direction system described in the fly’s brain, whereby sequences of visual or vestibular velocity inputs are integrated into a sequence of neural representations of the angular position of the animal. When training NLSC on this task, we showed that gate neurons segregated into two populations of neurons (Fig. 6c), as in hand-crafted models or as observed in the fly’s brain (5; 3). To understand the emergence of these two populations, we mapped the head-direction task onto a XOR problem (Fig. 5), and we followed the approach of (8) to interpret gate neurons as a hidden layer of neurons that non-linearly mix neural representations of the dictionary’s state and external inputs. We further argued that this segregation is a way to leverage sparse coding to build efficient neural representations that minimize interferences between the connectivity patterns programing transitions between states, given a fixed number of neurons.

In the head-direction case, external inputs are two dimensional, which is low compared to the number of dictionary states encoding angular position. We studied another extreme case where external inputs are high-dimensional compared to the number of dictionary states. In this case, we have shown it is more efficient to have gate neurons segregated in a number of populations that equals not the external inputs dimension but the number of dictionary states, with dictionary states modulating the gain of populations of gate neurons, and external inputs driving specific neural patterns in the gate neurons allowed to be active (Fig. 7c). Again, this segregation of gate neurons in optimized networks is a consequence of using sparse coding to minimize interferences among connectivity patterns. The intermediary case in which the number of external inputs and dictionary states are of the same order remains to be investigated.

### NLSC as a robust implementation of finite-state automata

We have shown that NLSC implements finite-state automata, with attractor-states of the dictionary implementing *P* machine states, and activation of gate neurons by *P*_*c*_ external inputs implementing transitions between machine states. Older works treated neural networks as automata. McCulloch and Pitts put forward a recipe to implement any finite-state automaton with a neural network ((26), Fig.3.5-1). Their implementation relies on an array of *P*_*c*_ × *P* neurons that are properly wired with each others as well as with sensors and effectors. Neural representations are not distributed, for instance activation of a machine state at a given time is supported by the activation of a single neuron. In NLSC, machine states are rather encoded by attractor-states, supported by recurrent connectivity, and transitions are programmed through distributed neural representations in gate neurons. An advantage of this FSA implementation is robustness, an important aspect regarding the implementation of computational devices in biological systems. Previous studies of perceptrons and attractor neural networks (42) have shown how to control the level of robustness by introducing a margin parameter that allows these systems to be robust to failure in synaptic release, neuronal loss or noisy inputs. Such a margin parameter could be added to our binary models, we expect the minimal number of neurons required to solve a given computational problem to increase smoothly with the margin value (42), without changing the scaling relationships between number of neurons and computational demand. Another advantage of the NLSC architecture regards these scaling relationships: leveraging sparse coding, the number of neurons required for implementing a FSA with a NLSC scales sub-linearly with the number of machine states or the number of external inputs, e.g. 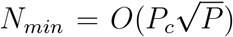. By contrast, the implementation of McCulloch and Pitts requires a network size that grows linearly with both *P* and *P*_*c*_. Recent work (45) has proposed an implementation of FSA using attractor neural networks, with direct hetero-associative connectivity to program transitions. They characterized the storage capacity of these systems through simulations, and also found that sparsely coded neural representations can be used to reduce network sizes.

Another important difference between NLSC and the neural networks proposed by McCulloch and Pitts regards the timing of computations. In their implementation, time is updated at each input presentation and all neurons are updated synchronously. Thanks to the stability of attractor states, NLSC can operate asynchronously (46), with a notion of time independent of the presentation of external inputs, even though these inputs still act as a clock to trigger transitions among machine states. In the last set of experiments we presented, we asked a NLSC implementing a sequential machine to autonomously update its state. This suggests that network oscillations (47) could serve as internally generated clocks, leading attractor states to become metastable (34).

### Relationship with brain circuits

In dictionaries of NLSC, neural dynamics exhibits attractor states which have been shown to be well suited to encode cognitive variables involved in various brain computations (48). By contrast, experimental evidence is scarcer for the gate neurons of the NLSC. The most direct evidence comes from studies of the head-direction system in flies, with the identification of two populations of protocerebral bridge neurons mixing clockwise or counter-clockwise vestibular velocity and current angular position (3). As detailed in the main text, our theoretical predictions do not match the low numbers of neurons observed in the fly’s brain, possibly due to some oversimplified assumptions of our modeling for the head-direction system of flies. Other studies with C. Elegans, focused on sequences of motor patterns, described a neuron that plays the role of a gate neuron, with its activation triggering transitions between different locomotive states (49; 50).

Beyond these evidence for gate neurons in invertebrate brains, could mammalian brain circuits implement NLSC ? As mentioned, cortical networks with their rich recurrent connectivity are well suited to implement dictionaries. A possible neural substrate for gate neurons is thalamic circuits, as proposed by previous modeling studies involving networks with gate-like neurons (44; 43). The cognitive variable interpretation of NLSC, in which gates allow for the reconfiguration of interactions between cognitive variables (SI Fig. 2, Method 7.4), assigns a well defined cognitive role for gate circuits. This is in line with the idea that cortical’s functional connectivity is under the control of thalamic activity (51). Recordings of thalamic activity in a delayed sensorymotor transformation task have shown that thalamic neurons respond to the go cue (16), it would be interesting to see whether thalamic neurons do so with patterns of activity that depend on the current cortical state. Our capacity calculations have shown that for NLSC implementing multiple permutations of dictionary states, the total number of gate neurons is of the same order or larger than the number of dictionary neurons. Given the relatively low number of thalamic neurons compared to neocortical ones (roughly one for ten), cortico-thalamic circuits do not appear well suited to store long-term engrams of NLSC. Future work will point at fully cortical circuits that could support the implementation of NLSC.

Whether or not transitions between neural states are orchestrated through specialized gate neurons, which could be associated with a specific cell class, remains to be determined. The characterization of NLSC we presented here allows to look for associated signatures in brain circuits.

We associated specific selectivity profiles with dictionary or gate neurons. In our trained ANN a dictionary neuron is associated to a single dictionary state. This results from the constraint of fixed read-out connectivity weights that we imposed to ease reverse-engineering. We have relaxed this assumption in binary neurons implementation of NLSC, where individual dictionary neurons exhibit mixed-selectivity for dictionary states, with a dictionary neuron participating in encoding multiple dictionary states. This is to be contrasted with the selectivity profiles of gate neurons which exhibit non-linear mixed-selectivity, responding to combinations of currently active dictionary state and external input. Gate neurons are expected to be segregated into populations, not necessarily spatially segregated anatomical structures as is the case for the head-direction system in flies, but at least in selectivity space (52; 18). Analysis methods of brain recordings have been proposed to define populations of neurons based on their selectivity (53; 54), they could be used to identify the predicted population structure of NLSC in the brain of animals performing specific behaviors. In particular, we expect to observe different population structures in gate neurons of NLSC expanding sequences of low-dimensional external inputs into a higher-dimensional space (*P* ≫ *P*_*c*_ for the head-direction system example, Fig. 6), compared to NLSC contracting high-dimensional external inputs into sequences of dictionary states lying in a lower-dimensional space (*P* ≪ *P*_*c*_ for the control of motor effectors example, Fig. 7). A distinction between these two architectures for NLSC lies in the arrangement of modulatory and driving pathways. Whether neurons are driven or modulated by their pre-synaptic inputs has been characterized experimentally in different brain structures (e.g. (27; 55)), such approaches could be used to identify NLSC in brain circuits.

When training ANN to balistically produce a sequence of neural states (Fig. 9), we observed that dictionary neurons, in addition to their selectivity to machine states, exhibit ramping up or ramping down activity, as observed in the motor-cortex analogue of singing birds (35) or in cortical networks during working memory tasks with delays with a characteristic time scale (36; 37; 38). In our models, such ramping controls the opening time of gate neurons (see also (56)), which differs from previous accounts of ramping activity (36). It would be interesting to manipulate the activity of populations of such ramping neurons to see whether it could elicit the activation of identified gate neurons.

Besides characterizing neural selectivity through coding levels or the population structure of gate neurons, the theory for optimized networks predicts different connection probabilities among dictionary and gate neurons. Gate neurons do not need to be connected with each others, this fits well with the low level of recurrence of thalamic circuits. Moreover, depending on the computation associated with a NLSC, the theory predicts different connection probabilities in between dictionary and gate neurons, or among dictionary neurons. Such signatures could be used to identify NLSC from connectomics data sets which provide connection probabilities between different cell classes (57; 58).

### Limitations and perspectives

In the NLSC we studied here, the dictionary is a Hopfield-like model with stable fixed-points that are randomly dispersed throughout state-space. For specific computations such as the path-integration implemented by the head-direction system, there is a natural notion of proximity between dictionary states encoding nearby positions. This has been traditionally modeled with the ring model, a particular instance of a continuous attractor neural network in which stable fixed-points lie on a manifold (59). Our quantitative characterization of NLSC could be extended to describe how the number of gate and dictionary neurons are reduced for continuous attractor neural networks or for networks with correlated patterns (60). This could allow to better relate NLSC to quantitative characterizations of the anatomy and physiology of the head-direction circuit (3; 61). A path-integration computation is also thought to involve grid-cells in medial enthorinal cortex (62). It would be interesting to describe the selectivity properties that gate neurons have in such a case, which would allow to propose different experimental predictions compared to alternative path-integration mechanisms (63; 62). Another example of brain structure that has been modeled with continuous attractors is area CA3 of the hippocampus. In this case, the dictionary is composed of multiple manifolds of fixed-points, each encoding a spatial map (39; 40). How path-integration could be performed for multiple maps in such models is an interesting theoretical problem, though it is not clear which inputs control the motion of bumps of activity in CA3.

In most of the situations we studied, external inputs trigger transitions between neural states. While this is fine for modeling systems computing on sensory inputs, it appears limited to model motor systems for which clocking signals have not been reported. In the last problem we studied, we tuned the connectivity of a NLSC, using gradient descent, so that it autonomously updates its active dictionary state. We have not characterized the connectivity changes that allow network states to become metastable (34). It would be interesting to understand how such a connectivity component would interfere with both the auto-associative and the hetero-associative components encoding the computation performed by a NLSC.

Throughout this article we have made observations about the selectivity and connectivity properties of trained artificial neural networks. Using binary neuron models we have shown how these properties allow to minimize the number of neurons. It remains to be understood why we observed such properties in artificial neural networks, which have been trained without any pressure on minimizing the number of neurons. A possibility is that those properties are also beneficial to overcome the noise we injected in these systems, or that the backpropagation algorithm we used enforce a form of regularization (64).

Studies of attractor neural networks have introduced order parameters to characterize the dynamics of collective patterns of neural activity (29; 65) and to model activations of the cell assemblies introduced by Hebb (15). This conceptual framework has proven useful to interpret neural recordings in behaving animals, with order parameters encoding cognitive variables, such as the identity of a remembered image (66), a location in space (67), a motor configuration (68), a choice option (69), or the integrated representation of a stream of sensory inputs (70). Cognition relies not only on these neural representations but also on the ability to switch between them. Here we have shown how external inputs, orthogonal to the dictionary’s manifold (71), orchestrate these switches by controlling the couplings between cognitive variables, through gain modulation in population-structured networks (18). Brain circuits receive inputs from external sensors but also and mainly from other distant brain areas. Also, elaborated cognitive processes presumably rely on multiple interacting circuits. Coupling attractor neural networks together has been shown to elicit rich temporal dynamics of the order parameters associated with each network (72; 9; 73). Future work will show how NLSC can be coupled with each others to support cognition, by running computations beyond those performed by finite-state automata (26).

## Acknowledgments

We are grateful to Gianluigi Mongillo for discussions and a careful reading of the manuscript.

## Author contributions

A.D. designed and performed research. A.L. discussed analysis of trained ANN, R.M. discussed analytical calculations. A.D. wrote the manuscript with inputs from all authors.

## Data and materials availability

All codes, for training ANN, solving analytical equations and generating figures, will be made available on a public repository at the time of publication.

## 7 Methods

### 7.1 Artificial neural networks

We trained networks of rate units that are discretized version of the continuous time equations

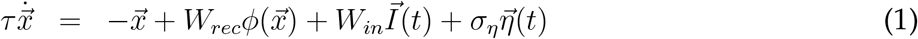

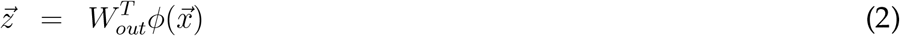

where 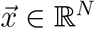 is the vector of currents to the *N* neurons composing the network. Currents are injected in the network by *N*_*in*_ input neurons 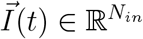 through *N*_*in*_ input weight vectors collected in the matrix *W*_*in*_ ∈ ℳ (*N*_*in*_, *N*). We divide inputs into two categories 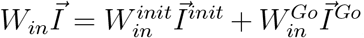, where 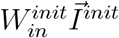 stands for input used to initialize networks into specific stable states and 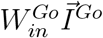 for inputs triggering transitions between stable states. Currents flow in the network through the recurrent matrix *W*_*rec*_ ∈ ℳ (*N, N*). These currents are transformed into firing rates through the activation function *ϕ*(.) which we take to be the rectified linear unit except for low-rank networks for which we used the hyperbolic tangent that leads to more interpretable mean-field equations. *N*_*out*_ output neurons 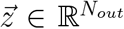 read linear projections of the firing rate vector 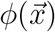 through *N*_*out*_ output weight vectors collected in the matrix *W*_*out*_ ∈ ℳ (*N, N*_*out*_). 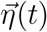 is a vector of independent white noises, with standard deviations *σ*_*η*_ = 0.05.

Training is performed using the Pytorch library by tuning specified connectivity parameters so as to minimize a loss 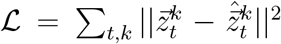, where *k* indexes trials in the input-output dataset and *t* indexes discretized time (we used a discretization time step of 20ms, for a time constant *τ* = 100ms). 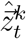 represents the desired output activations, while 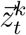 represents the actual output activations. Minimization is performed with stochastic gradient descent using the Adam optimizer with standard parameters. The choice of connectivity parameters used to minimize the loss depends on the specific training set-ups we used.

### 7.2 Training set-ups

#### 7.2.1 Training NLSC

Networks are segregated into *N* = *N*_*dic*_ + *N*_*gate*_ neurons through constraints on the connectivity (SI Fig. 1a). The *N*_*dic*_ neurons receive inputs from the initializing inputs 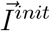 and are connected to the readouts, while the *N*_*gate*_ neurons receive inputs from the transition inputs 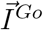 and do not connect to the readouts. The *N*_*dic*_ dictionary neurons are recurrently connected with each others, while the *N*_*gate*_ gate neurons are not, though gate neurons are bidirectionally coupled with dictionary neurons. Dictionary connectivity parameters 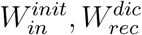 are trained on the dictionary task to implement *P* stable states (Method 7.3.1) while output connectivity *W*_*out*_ are fixed with readout neurons receiving from *P* non-overlapping subsets of dictionary neurons to ease interpretation of trained networks. Input connectivity to gate 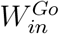 and gate/dictionary connectivities *W*^*dg*^, *W*^*gd*^ are trained to implement task-specific transitions between dictionary states. All initial connectivity parameters are drawn from Gaussian distributions. The time constant of gate neurons is decreased from 100 to 20ms to obtain sharper state transitions. Except when notified, *N*_*dic*_ is equal to *P* × 41 and the number of gate neurons equals the number of dictionary neurons.

#### 7.2.2 Training unconstrained networks

Neurons are not segregated into dictionary and gate neurons, and we train an unconstrained recurrent matrix *W*_*rec*_ (SI Fig. 1b) to implement stable states and task-specific transitions. Input weights 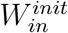 and output weights *W*_*out*_ are fixed to values obtained by independently training a network on the dictionary task, while transition inputs 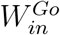 target all neurons with a fixed connectivity weight drawn from a centered normalized Gaussian distribution.

#### 7.2.3 Training low-rank networks

Recurrent matrices of rank-K are parametrized as 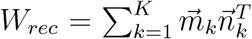 with 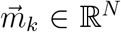. The rank K corresponds to the number of dictionary states, and connectivity vectors 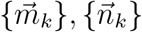 are initialized as Gaussian vectors structured into populations ((18), SI 8.2) to instantiate the relevant neural architectures. Each population is composed of 512 neurons. Values of the connectivity vectors are then fine-tuned to perform the task of interest by performing gradient descent on the loss parametrized with the 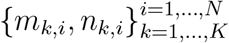 instead of the {*W*_*ij*_}_*i,j*=1,…,*N*_.

### 7.3 Descriptions of individual tasks

In all tasks, networks are first trained on trials corresponding to individual transitions making up the behavior of interest. These trials have a fixation epochs of duration *T*_*fix*_ = 100ms, followed by a *cueing epoch* of duration *T*_*cue*_ = 100ms where the dictionary is set into its initial state through activation of input neurons 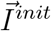. It is followed by a *maintenance epoch* of durations randomly drawn at each trial *T*_*main*._ ∈ [500, 2500]ms and a *go epoch* of duration *T*_*go*_ = 200ms during which the go signal is applied to the network 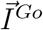, followed by a *response epoch* of duration *T*_*resp*_ = 1000ms.

#### 7.3.1 Dictionary task

In order to build a dictionary of *P* states, we define an initializing input vector 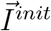, and a readout vector 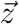 of size *P*. A single input neurons *j* ∈ [1, …, *P*] is activated at a value 1 during the *cueing epoch* if the trial starts with maintaining dictionary state *X*_*j*_. We set the target activations 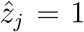 and 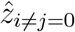. Here, and only for this task, we do not introduce additional transition inputs 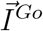, rather initializing input neurons 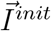 can also be activated during the *go epoch*. If input neuron *k* ∈ [1, …, *P*] is activated at 1 during the go epoch, we trained networks with target output activations 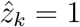 and 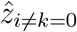, while if the input neuron is activated at 0.5, 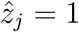 and 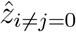 (SI Fig. 1c).

Networks trained on this task develop positive feedback loops among neurons belonging to the same cell assembly, defined as neurons projecting to the same readout neuron, as well as inhibition among neurons belonging to different cell assemblies. This connectivity structure leads to attractor dynamics making dictionary states robust stable states (**?**).

#### 7.3.2 Sensory-motor transformation task

Two initializing inputs 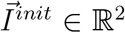 initialize the network in either state *A* or *C*. Upon activation of the single go input neuron *I*^*Go*^, the network is asked to transit to state *B* or *D* (SI Fig. 3a).

We trained NLSC on this task. Their dictionary network is characterized by positive feedback loops among cell assemblies associated to each dictionary state (SI Fig. 3b). With such a connectivity, recurrent currents generated by the maintenance of state *A* (or *C*) are similar onto neurons of state *B* and state *D* (SI Fig. 3c right-top). By contrast, specific patterns of gate neuron activation during the *go epoch* (Fig. 3c left) produce asymmetric currents onto cell assemblies *B* and *D* (SI Fig. 3c, right bottom), triggering a transition towards the correct next cell assembly.

We also trained 50 unconstrained networks on this task. They share similar characteristics to those of the NLSC, with positive feedback loops among cell assemblies associated to each dictionary state (SI Fig. 3d). Neurons belonging to cell assemblies *C* or *D* playing the role of gate neurons during transitions from *A* to *B* (SI Fig. 3e, left) and sending transient asymmetric currents to cell assemblies *B* and *D* (SI Fig. 3e, right bottom). In addition to this source of asymmetry, inhibitory connectivity between neurons belonging to different cell assemblies (SI Fig. 3d) are asymmetric, contributing to the difference in currents to cell assemblies *B* and *D* at the time of transition (SI Fig. 3e, right top).

In fact, for this specific task, simply shaping the direct connectivity among cell assemblies is enough, without the need to resort to gate neurons. This is shown by the network of SI Fig. 3f whose connectivity is initialized by adding strong inhibition from cell assembly *A* to *D* and from *C* to *B*, and which performs the task without gate neurons (Fig. 3g left). In this case, the symmetric go cue provides the same level of excitation onto cell assemblies *B* and *D* (SI Fig. 3g right bottom), but in trial *A, B* is less inhibited than *D* throughout the maintenance epoch (SI Fig. 3g right top) and thus *B* activates.

Training a rank 4 network on the task allows to get the compact descriptions of the recurrent dynamics of Fig. 1f. It is characterized by 4 cognitive variables that take non-zero values at specific times (SI Fig. 3h), with functional connectivities between them that are controlled by the go cue (SI Fig. 3i) enabling neural activity to flow from active cell assembly *A* to *C* or from *B* to *D*.

#### 7.3.3 Single permutation of dictionary states

*P* = 20 initializing input neurons 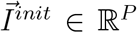 send input to the networks, with a single of them *k*_0_ ∈ [1, …, *P*] activated to 1 during the *cueing epoch*, instructing output neuron 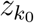 to be active during the *maintenance epoch*. Upon activation of the single go input neuron *I*^*Go*^, the network is asked to transit from state *X* to state *X* + 1 as reported by activation of the output neurons 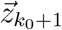 (Fig. 4a).

We trained NLSC on this task and reported their stereotyped behavior through an example network. For each transition a gate activation pattern is obtained by averaging the firing rate of each gate neuron over the *go epoch* (red vertical lines in SI Fig. 4a bottom). The distribution of firing rate of gate neurons at transitions is reported in SI Fig. 4b. To report statistics on the patterns of activations of gate neurons in the main text, we defined a gate neuron to be active in a transition if its firing rate is larger than 0.05. Examining pairs of transitions reveals they are associated with orthogonal gate activation patterns as shown in SI Fig. 4c. In the NLSC implementations, training the dictionary on the dictionary task leads to positive feedback loops among neurons belonging to the same cell assembly, with inhibition between neurons belonging to different cell assemblies. As expected from this training set-up, the inhibition is symmetric and independent on the transition path, i.e. state *X* does not inhibit less state *X* + 1 compared to other states (SI Fig. 4d). As a result, currents coming from a cell assembly activated during the *maintenance epoch* are self-exciting and equally inhibit all other states (SI Fig. 4e right top). By contrast, currents produced by activation of the gate patterns send excitatory inputs to dictionary neurons belonging to the next cell assembly to be activated (SI Fig. 4e right bottom).

We also trained 50 unconstrained networks on this task. They develop positive-feedback loops among neurons activated in a same cell assembly (SI Fig. 4f). An asymmetry in inhibitory connectivity is observed, with less inhibitory connectivity from neurons of cell assembly *X* to neurons of cell assembly *X* + 1 compared to the other (SI Fig. 4f). This leads currents produced by the maintenance state to provide *slightly* less inhibition to neurons belonging to the next cell assembly to be activated (SI Fig. 4g right top). At transition time, neurons associated neither with the maintenance state nor the response state are activated (SI Fig. 4g left), leading to provide *largely* more excitatory currents onto neurons of the response state (78% of the asymmetry in currents, SI Fig. 4g right bottom). In all the trained networks transient activations during the *go epoch* are necessary to trigger the transitions from one cell assembly activation to another. This is shown in SI Fig. 4h where we report that selectively inactivating the 50 most transiently active cells at transition *k*_0_ disrupts this transition but not the others.

The switching currents arise mainly from gate-like neurons in this scenario, as opposed to the case of the sensory-motor transformation task (Fig. 1h and Fig. 2b). We note that networks for the permutation task, involving 20 dictionary states, are larger (20*41 instead of 4*41 neurons). We checked whether the observed differences in switching currents could be due to network size, as there are more neurons that can develop gate-like properties, rather than number of transitions as argued in the main text. To do so, we trained 15 large ANN with 20*41 dictionary neurons on the sensory-motor transformation task, and found that, as for smaller networks, switching currents remained dominated by direct currents (80%/20% of the asymmetry in switching currents from direct/gated currents).

#### 7.3.4 Head-direction system and multiple permutations of dictionary states

Here we consider a task where networks are trained to switch between *P* states according to *P*_*c*_ > 1 permutations of the *P* states. To model the head-direction system, we considered the case *P*_*c*_ = 2, with the two permutations being the +1 shift and the −1 shift once the *P* = 20 states are ordered on a ring (SI Fig. 5a). *P* initializing inputs 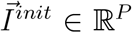 send input to the networks, with a single input neuron *k*_0_ ∈ [1, …, *P*] activating to 1 during the *cueing epoch*, instructing output neuron 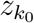 to be active during the *maintenance epoch*. Two inputs can trigger transitions 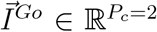. Upon activation of one of this input neurons during the *go epoch*, the network is asked to transit from state *X* either to state *X* + 1 as reported by activation of the output neurons 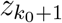, either to state *X* − 1 as reported by activation of the output neurons 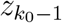. In this case, the number of permutations is much lower than the number of states *P*_*c*_ ≪ *P*.

We also trained networks on a task where *P*_*c*_ ≫ *P*, implementing all the possible *P*_*c*_ = 4! = 24 permutations of a set of *P* = 4 dictionary states (SI Fig. 5h).

We trained NLSC on the first task and report their stereotyped behavior through an example network. For all the 2 × 20 transitions, we analyzed gate activation patterns that can be visualized in the bottom part of the raster in SI Fig. 5b. To understand their organization we note that the trained connectivity 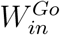 composed of two vectors gets structured upon training, with some gate neurons being preferentially targeted by one of the two transition inputs in 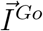, SI Fig. 5c top. Reordering gate neurons by splitting them into two groups, according to whether they receive more inputs from the clockwise or counter-clockwise input, reveals the structure of the neural activity patterns of gate neurons (SI Fig. 5c bottom). We confirmed that this structure plays a functional role by performing inactivation experiments: inactivating the first population of gate neurons during the *go epoch* prevents clockwise transitions (SI Fig. 5d top) and leaves unaffected counter-clockwise transitions (SI Fig. 5d bottom), as summarized in SI Fig. 5e.

We also trained 50 unconstrained networks on this task. They develop positive-feedback loops among neurons activated in a same cell assembly. As expected no asymmetry in inhibitory connectivity is observed from state *X* to states *X* +1 and *X* − 1, although these states are less inhibited than the others (SI Fig. 5f). This leads currents produced by the maintenance state (SI Fig. 5g left) to provide the same amount of inhibition to neurons belonging to the cell assemblies associated with *X* + 1 and *X* − 1 (SI Fig. 5g right top). At transition time, a pattern of neurons associated with other states (Fig. 5g left), specific to the activated go cue, decides the following cell assembly to be activated (SI Fig. 5g right bottom).

We trained NLSC on the second task with *P*_*c*_ ≫ *P* (SI Fig. 5h) and report their stereotyped behavior through an example network. We take *N*_*dic*_ = 41 ∗ 4 and *N*_*gate*_ = 1640. To obtain successful training, we scaled-up initial inputs 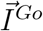 and scaled-down input vectors 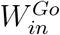 by a factor 100, allowing to balance learning speeds in *W*^*dg*^ and 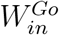. In order to understand how training shapes neural activity in the gate neurons (SI Fig. 5i bottom), we examined the average *in-going* connection strength to individual gate neurons, from dictionary neurons associated to one of the *P* = 4 states. SI Fig. 5j top shows that this connectivity gets structured during training, such that each gate neuron could be meaningfully assigned to one of the dictionary state. This is done by grouping the *N*_*gate*_ gate neurons into *P* = 4 clusters, using a Gaussian mixture clustering algorithm (SI Fig. 5j bottom). Reordering the X-axis of the raster of gate patterns according to these four groups, as well as the Y-axis, grouping transitions originating from the same dictionary states, reveals the organization of gate patterns (SI Fig. 5k top). In comparison, reordering the raster after performing clustering on averaged *out-going* connection strengths from individual gate neurons to dictionary neurons associated to one of the *P* = 4 states, did not show any sign of meaningful structure (SI Fig. 5k bottom). We confirmed that the meaningful structure obtained by clustering plays a functional role by performing inactivation experiments: inactivating, during the *go epoch*, the population of gate neurons associated to state *X* disrupts transitions where *X* is a maintenance state but not the other transitions (SI Fig. 5l).

#### 7.3.5 Motor sequence without go cue

Here we trained NLSC on the same permutation task as described in Method 7.3.3, except that there is no external go cue instructing the network to transit. Instead, the loss function is such that the network is asked to maintain each state for a predefined duration *T* = 250, 500, 750, 1000, 1500ms. The power spectrum of the average activity of gate neurons (Fig. 9c) is obtained using the *welch* algorithm of the *SciPy* python package with the *hann* window. In order to obtain Fig. 9d,e,f we perform a PCA analysis aligning individual dictionary neurons on a single temporal window of duration *T′* = 1.2*T*. Instead of looking for population modes as traditionally done, we defined temporal modes by performing PCA on a correlation matrix of size 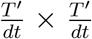, obtained from the aligned raster of dictionary neurons activation.

### 7.4 Mean-field theory for low-rank networks

In order to visualize neural activity as trajectories in state space for the network implementing the sensory-motor task (SI Fig. 2), we built a low-rank network with Gaussian vectors segregated into populations (18). Its dynamical behavior **(1)** can be reduced to interactions between cognitive variables, which characterize the location of neural activity along directions 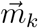:

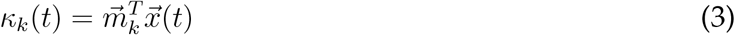

The dynamics is reduced to a dynamics over four variables, with a specific form obtained by shaping the connectivity matrix (SI Section 8.2):

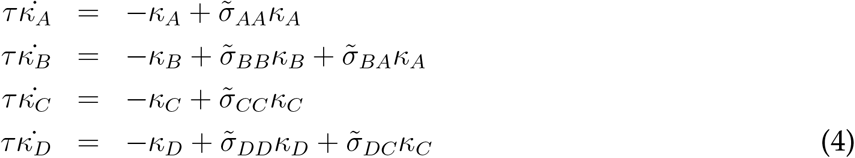

where the functional connectivities 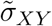 are non-linear functions of the four cognitive variables and the go cue input, these functions are parametrized by connectivity’s summary statistics. In particular the go cue input, by setting the gain of neurons in the gate population from 0 to 1 (SI Fig. 2d), is able to control the values of 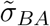 and 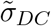 (SI Fig. 2b) and to trigger the transitions from the state (*κ*_*A*_ = *c, κ*_*B*_ = 0, *κ*_*C*_ = 0, *κ*_*D*_ = 0) to the state (*κ*_*A*_ = 0, *κ*_*B*_ = *c, κ*_*C*_ = 0, *κ*_*C*_ = 0) and from (*κ*_*A*_ = 0, *κ*_*B*_ = 0, *κ*_*C*_ = *c, κ*_*D*_ = 0) to (*κ*_*A*_ = 0, *κ*_*B*_ = 0, *κ*_*C*_ = 0, *κ*_*D*_ = *c*) (SI Fig. 2a), with *c* = *O*(1) a constant. The flow fields plotted in SI Fig. 2c represent the time evolution of cognitive variables 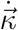 obtained from e.g. equation **(4)**, each panel is obtained using the *streamplot()* function of *matplotlib* on 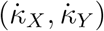.

### 7.5 Capacity of neural architectures

Our results on the minimal number of neurons, *N*_*min*_, required to implement NLSC-based architectures build on previous results obtained for perceptrons with binary neurons and binary synaptic weights (20; 21; 74; 22). We consider *N*_*out*_ perceptrons with *N*_*in*_ input neurons. The *P*_*a*_ input patterns 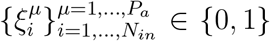 are drawn randomly with the constraint 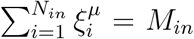 and output activations 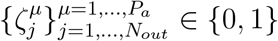 are drawn randomly.

We present calculations for computing the probability that the synaptic connectivity matrix 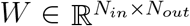 is not overloaded by the stored associations, i.e. that the output neurons correctly classify the *P*_*a*_ input patterns: 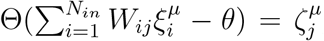, for all *µ* = 1, …, *P*_*a*_ and *j* = 1, …, *N*_*out*_, with Θ(.) the Heaviside function and 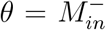 the activation threshold. The matrix of binary synaptic weights is assumed to undergo Hebbian learning while the *P*_*a*_ correct input and output activations are imposed onto the network: 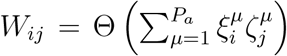. We note *g* the fraction of synapses activated after presenting *P*_*a*_ associations. It is a crucial quantity for our analysis as it controls the amount of interferences between associations and indicates for which value of *P*_*a*_ the learning rule is not able to correctly store the associations anymore. It can be estimated as

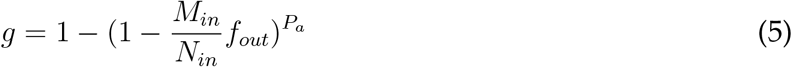

The probability that *P*_*a*_ associations in *N*_*out*_ perceptrons are learnt without errors can be expressed as (SI Section 8.1.1):

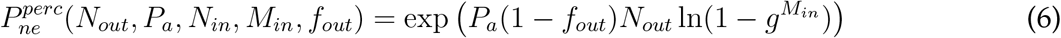

The neural architectures we built can be decomposed into multiple independent such perceptrons (Fig. 3a, SI Fig. 7a-d). The probability that all the associations of the architecture are learnt without errors, 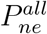, can thus be estimated by a product of the functions **(6)** estimated with the appropriate input variables. In order to estimate *N*_*min*_ for a given architecture and given numbers of associations, we performed grid-search over the number of neurons and coding properties *N*_*in*_, *M*_*in*_, *f*_*out*_ and look for sets of parameters that minimize the total number of neurons while keeping the probability of no error above a threshold 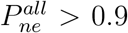. For instance, an attractor neural network of *N* neurons can be seen as *N* perceptrons (SI Fig. 7b). For *M* active neurons per pattern and *P* patterns, we compute 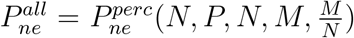 and perform grid-search on parameters *N* and *M* to find the values that minimize *N* keeping 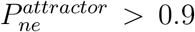. In SI Section 8.1, we detail this process for each of the neural architectures considered in the main text.

## 8 Supplementary

### 8.1 Capacity calculations

#### 8.1.1 Capacity for perceptrons

We detail how to obtain the probability **(6)** that the *P*_*a*_ associations are perfectly retrieved for *N*_*out*_ perceptrons with input size *N*_*in*_, input coding level 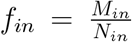 and output coding level *f*_*out*_ (SI Fig.7a). We do so following (22). To simplify mathematical expressions we have considered the case where the number of active neurons in the input patterns is fixed and does not fluctuate from pattern to pattern: 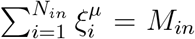 for all *µ*’s. With this constraint, the activation threshold of the output neurons can be taken as 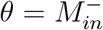, and, given the learning rule leading to

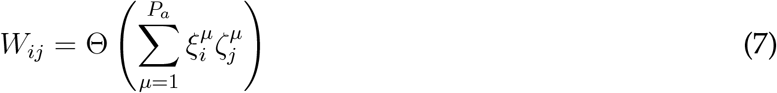

we are guaranteed that no errors will be produced when retrieving the *P*_*a*_*f*_*out*_ associations with 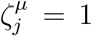. Errors can then only occur for associations with 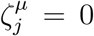. They occur when the output neuron receives an input *h* = *M*_*in*_, i.e. when all the synapses between the output neuron and active input neurons in 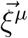 have been potentiated by one of the other *P*_*a*_ − 1 associations. The probability of such an event can be simply estimated by

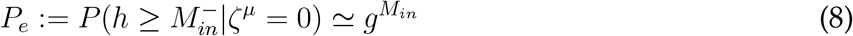

with *g* taken from **(5)**. The probability that no errors are produced in all the *P*_*a*_ associations and *N*_*out*_ perceptrons is then given by

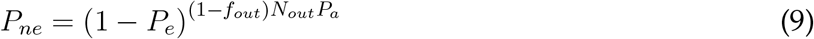

for sparse enough coding levels 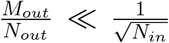 correlations between two synapses 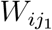 and 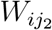 can be ignored (25) and the probability of error is well estimated by **(8)**. In order to obtain more precise estimates for finite-size networks and compare with simulations (Fig. 3b, crosses) we took these correlations into account using exact calculations recapitulated in (75) equation (42):

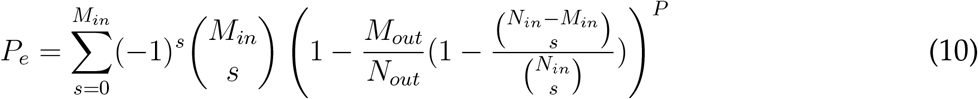

This expression is cumbersome to estimate, such that when optimizing architectures made of multiple NLSC we used **(8)** instead. SI Fig. 7h shows that the two estimates of *N*_*min*_ are very close to each other and follow the same dependency on parameters. Only the parameters *M*_*in*_ and *M*_*out*_ at which *N*_*min*_ is found are slightly different, which can be taken into account to improve the match with simulations.

#### 8.1.2 Storage capacity of attractor neural networks

Here we apply the above results to compute the minimal number of neurons *N* required to store *P* fixed-points in an attractor neural network. This problem is equivalent to storing *P* associations in *N* perceptrons with input layer of size *N* and equal input and output coding levels 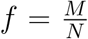 (SI Fig. 7b). Applying equation **(6)** to our problem, we can estimate the probability of no error

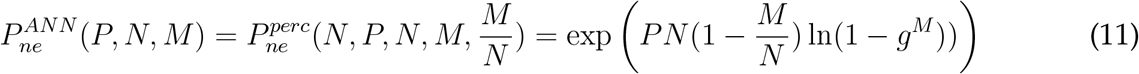

In figure Fig.3b dashed line, we show how the minimal number of neurons N increases when more fixed-points are to be stored. It is obtained by computing 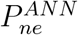 for a region of parameters (*N, M*) and report the minimal value of *N* such that *P*_*ne*_ > 0.9. Crosses represent the minimal value of N obtained with simulations, they are obtained as follows: we built 100 attractor neural networks with parameters *N, M, P*, we tested the stability of all the *P* patterns for each, and reported the probability that no error occurs for all the P patterns *P*_*ne*_ ; for each cross we fixed *P* as well as the value of *M* that is found to optimize the capacity according to **(11)**, then simulated networks with various values of *N* and reported the smallest value for which *P*_*ne*_ > 0.9. As expected from the infinite-size estimate (22), the minimal number of neurons required to store *P* patterns of activity scales as 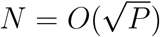.

#### 8.1.3 Capacity for the implementation of transitions between dictionary states

We consider a dictionary network of *N*_*dic*_ neurons storing *P* fixed-points 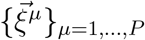 with coding level 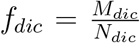, together with *N*_*gate*_ gate neurons (SI Fig.7c). We want this circuit to implement a total of *T* = *P* transitions between the fixed-points of the dictionary with each transition associated to a pattern of activation of gate neurons 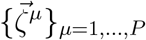 with coding level 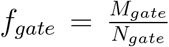. To trigger a transition, the appearance of a go cue at time *t* has the following two effects: a strong inhibition on gate neurons is released, so that at time *t* + 1 gate neurons represent pattern 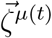 if the dictionary was in pattern 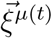, as well as an increase of the activation threshold in the dictionary, so that the dictionary forgets its current pattern. With this, at time *t* + 2 the dictionary is put into 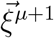 by the pattern printed in gate neurons ^1^. As explained in the main text, constraints on such an architecture comes from i) storing the *P* fixed-points in the dictionary, ii) storing *P* associations from dictionary to gate neurons, and iii) from gates to dictionary neurons. Learning these associations under the Hebbian learning rule for binary synapses **(7)** leads to a fraction of potentiated synapses from dictionary to gates and gates to dictionary:

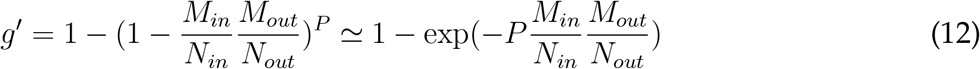

as in the previous section successful and efficient storage of the *P* associations is obtained for 0 < *g′* < 1. Applying **(6)** and **(11)** to i) the *N*_*dic*_ perceptrons of input-size *N*_*dic*_ of the dictionary, ii) the *N*_*gate*_ perceptrons of input-size *N*_*dic*_ from dictionary to gate, iii) the *N*_*dic*_ perceptrons of inputsize *N*_*gate*_ from gate to dictionary we get

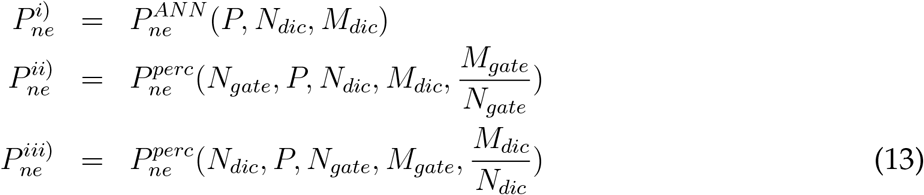

For a given value of *P* one can then scan parameters *N*_*dic*_, *M*_*dic*_, *N*_*gate*_, *M*_*gate*_ to find values that minimize the total number of neurons *N* + *N* under the constraint that 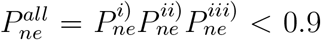 (estimates do not dramatically depends on the chosen threshold, not shown).

The full curve in Fig.3b shows the minimal total number of neurons *N*_*min*_ for storing *P* patterns as fixed-points of the dynamics and implementing a permutation, i.e. *P* transitions, among these fixed-points. SI Fig.7e shows variations with *P* of the number of dictionary and gate neurons, the coding levels in the dictionary 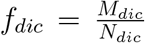 and gate 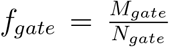 layers, and the fraction of activated synapses among dictionary neurons and in between gate and dictionary neurons.

In the set-up considered above specific patterns of activation of gate neurons 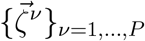 are learnt via Hebbian learning of the dictionary to gate connectivity. Instead random gate patterns could be implemented with a random binary connectivity, with inhibitory interactions among gate neurons tuning their coding level at a desired value 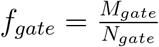. With such an assumption, constraint ii) on the dictionary-to-gate connectivity does not have to be taken into account anymore. Considering only constraints i) and iii), i.e. ignoring the second line in **(13)** leads to the orange curve in SI Fig. 7g.

If only a subset of dictionary states are associated with a transition *T* = *ϵP* < *P*, the number of necessary gate neurons is reduced linearly as shown in SI Fig. 7f obtained by replacing *P* by *T* to estimate the second and third lines of **(13)**.

#### 8.1.4 Capacity for permutations with direct hetero-associative connectivity

Most sequence models use direct hetero-associative connectivity (e.g. (76)) whereby neurons active in pattern *µ* directly excite neurons active in pattern *µ* + 1. In order to compare the performance of the NLSC with such a neural architecture, we consider a model of *N* neurons, with *M* active neurons per patterns, connected through

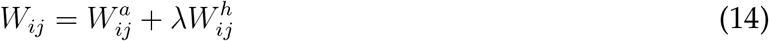

with 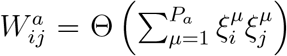 the auto-associative connectivity stabilizing neural patterns 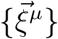 and 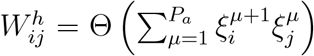 the hetero-associative connectivity directing transitions from neural pattern 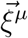 to 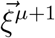. Compared to the NLSC, we do not have the extra gate neurons, but we expect more interferences due to the fact that both auto- and hetero-associative connectivity can activate neurons that are supposed to remain silent. As for the NLSC model, we take an activation threshold *θ* = *M* ^−^ when probing the stability of a neural pattern. Transitions are triggered by an external input *I*_0_, uniform on all neurons, and we take an activation threshold *θ* = (*I*_0_ + *λM*)^−^ when probing errors in transitions. With this, active neurons in 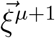 are sure to be activated. Activation of 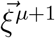 silences 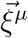 by a mechanism that we do not model. The probability of no error for this model is computed as 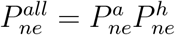 with

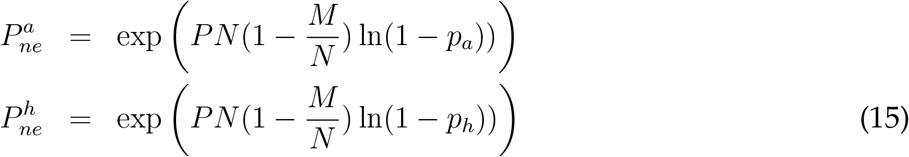

with

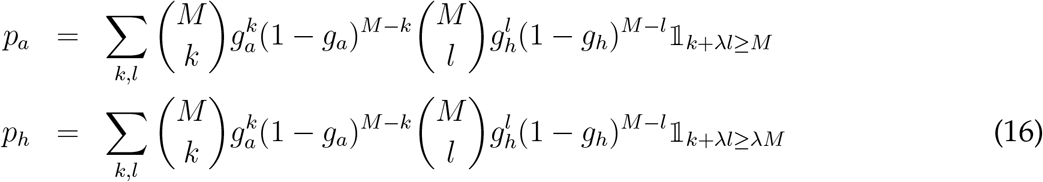

with *g*_*a*_ = *g*_*h*_ the fraction of activated auto-associative and hetero-associative synapses potentiated at 1. Fig. 3c shows that *N*_*min*_ is higher for all values of *λ* compared to the NLSC implementation.

#### 8.1.5 Capacity for multiple permutations of dictionary states

Instead of a single go cue associated with a single permutation, here we consider *P*_*c*_ go cues, or contextual signals, associated with *P*_*c*_ permutations of *P* dictionary states, *P*_*c*_ = 2 for the head-direction system. The role of gate neurons is to demix its *P*_*c*_ × *P* possible inputs (*P* configurations of the dictionary, *P*_*c*_ configurations of the go cue inputs) into linearly separable representations. This allows to learn the gate to dictionary connectivity so that a gate pattern elicit the correct next dictionary pattern. Training ANN in the different regimes *P*_*c*_ ≪ *P* and *P*_*c*_ ≫*P* revealed that patterns of gate neurons organize differently in these two extreme cases. In the first regime, a go cue input activates the gain (releasing inhibition brings neuron from below up to firing threshold) of gate neurons in one of *P*_*c*_ populations. Each dictionary pattern instead drives a pattern of activation of gate neurons in this sub-population (Fig. 6f). In the second regime, the situation is reversed, with dictionary patterns defining *P* populations of gate neurons by gain modulation (Fig. 7c), and the *P*_*c*_ go cue inputs driving patterns of activity in the activated sub-population. These arrangements can be understood as optimal from the capacity point of view. Indeed, the total number of gate neurons scales linearly with the number of populations, while it only scales with the square root of the number of driving patterns, thanks to sparse coding. The split into *P*_*c*_ populations thus gives 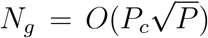 for *P*_*c*_ ≪ *P*, while the split into *P* populations gives 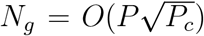. It remains to be seen whether there is a better way that can be proposed to mix dictionary and go cue inputs in the regime *P* ≃ *P*_*c*_.

In the regime *P*_*c*_ ≪ *P* with *P*_*c*_ populations of gate neurons, following the reasoning of the section above, the probability that there is no error for all the *P* stable states and all the *P*_*c*_ × *ϵP* transitions can be written as

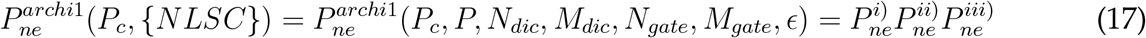

with

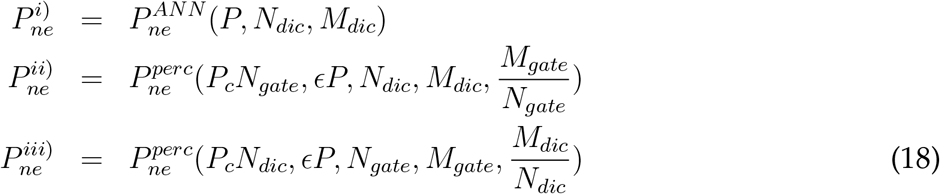

{*NLSC*} = {*P, N*_*dic*_, *M*_*dic*_, *N*_*gate*_, *M*_*gate*_, *ϵ*} contains all the parameters defining anatomical and physiological properties of the NLSC. Compared to the previous case **(13)** we simply took into account the fact that the number of perceptrons from dictionary to gate and gate to dictionary are proportional to *P*_*c*_. *N*_*gate*_ and *M*_*gate*_ here refer to the parameters of a single gate population, as is the case afterwards. To obtain *N*_*min*_ we looked for parameters that minimize the total number of neurons *N*_*min*_ = *N*_*dic*_ + *P*_*c*_*N*_*gate*_ while keeping *P*_*ne*_ > 0.9.

For the regime *P*_*c*_ ≫*P* all the *P* ^2^ possible transitions are wired into the network. To do so, all the go cues associated with the same transition drive the same pattern of neural activity in the activated population of gate neurons. This is done without any error since here neural patterns associated to each permutation are non-distributed, i.e. a single go cue neuron is activated in only one permutation, thus there are no interferences between go cue to gate neuron synaptic wires associated with different permutations. This explains the independence of *N*_*min*_ on *P*_*c*_ for this architecture (Fig. 8a, dashed line). Again the probability that there is no error for all the *P* stable states and all the *P*_*c*_ × *P* transitions can be written as

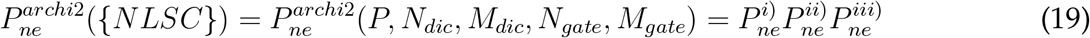

with

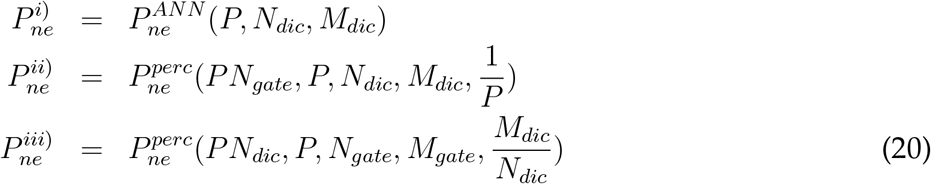

The second line is obtained considering *P* associations to be learnt on *N*_*out*_ = *P* ∗ *N*_*gate*_ perceptrons, with *f*_*in*_*N*_*in*_ = *M*_*dic*_ and an output coding level 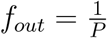. To obtain *N*_*min*_ we looked for parameters that minimize the total number of neurons *N*_*min*_ = *N*_*dic*_+*PN*_*gate*_ while keeping 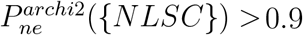.

### 8.2 Low-rank networks

We trained low-rank networks to perform the sensory-motor transformation task of Fig. 1. To do so we initialized the connectivity matrix as 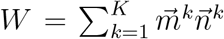 such that recurrently generated activity lies in a subspace of dimension *K* (77) and minimized the loss by updating the parameters 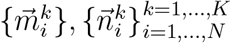, where *N* = *N*_*dic*_ + *N*_*gate*_ is the total number of neurons. Connectivity vectors are structured into populations (18) using the support vectors 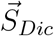 and 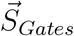 with non-zero entries at 1 on dictionary and gate neurons. We used the following rank *K* = 4 connectivity matrix associated with the four cognitive variables:

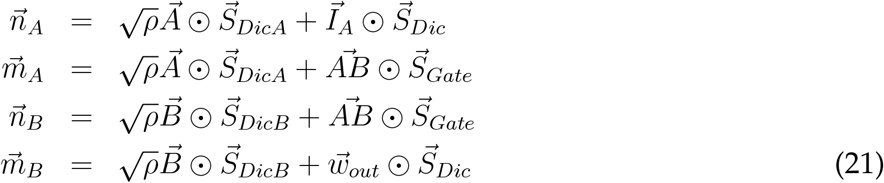

and symmetrically

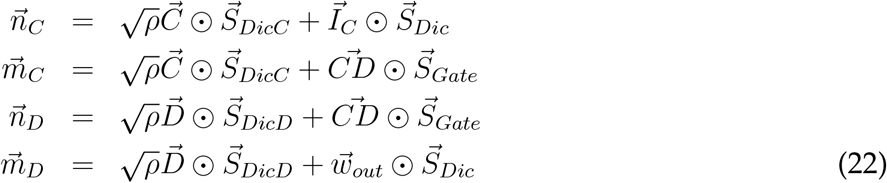

Vectors 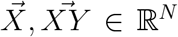 are Gaussian vectors with entries drawn independently from the normalized centered Gaussian distribution. The dictionary population is split into four populations with support vectors 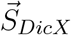 having non-zero entries at 1 on dictionary neurons of population X, so has to have four independent fixed points by taking *ρ* > 1 such that 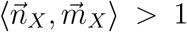. ⊙ is the Hadamard product performing entry-wise multiplication. Vectors 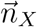 decide which inputs are picked up by recurrent activity. In order for the two cognitive variables *κ*_*A*_ and *κ*_*C*_ to activate in response to the two stimuli, we take 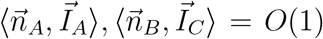, with 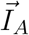 and 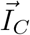 Gaussian sensory input vectors. Vectors 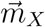 decide in which direction recurrent currents are generated. We take 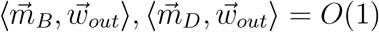 for generating the output, and 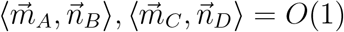 to transfer activations of *κ*_*A*_ or *κ*_*C*_ into activation of *κ*_*B*_ or *κ*_*D*_ upon presentation of the go cue, which changes the gain of gate neurons from 0 to 1 through the input (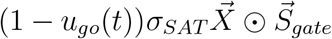 with *σ*_*SAT*_ ≫1.

Equations over cognitive variables 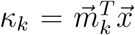 are obtained by projecting the dynamical equation **(1)** onto directions *m* _*k*_ leading to equation **(4)** (18) with

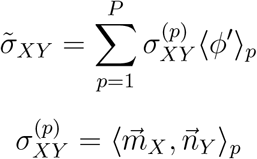

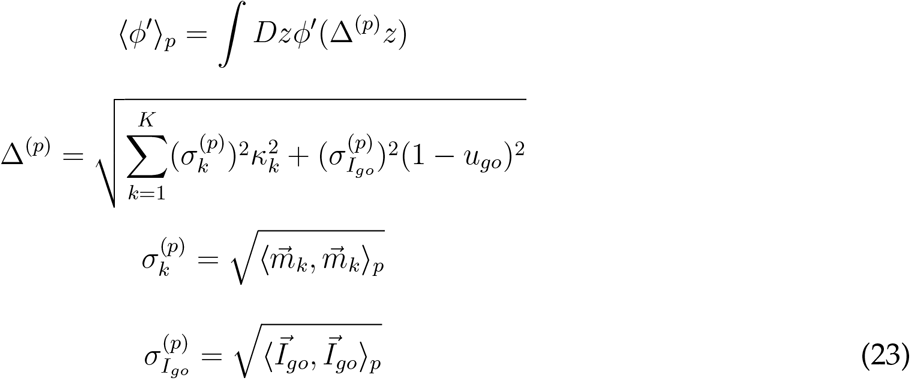

where the scalar products ⟨.,. ⟩ _*p*_ are taken over neurons belonging to population *p*.

## Supplementary figures

**SI Fig. 1:**
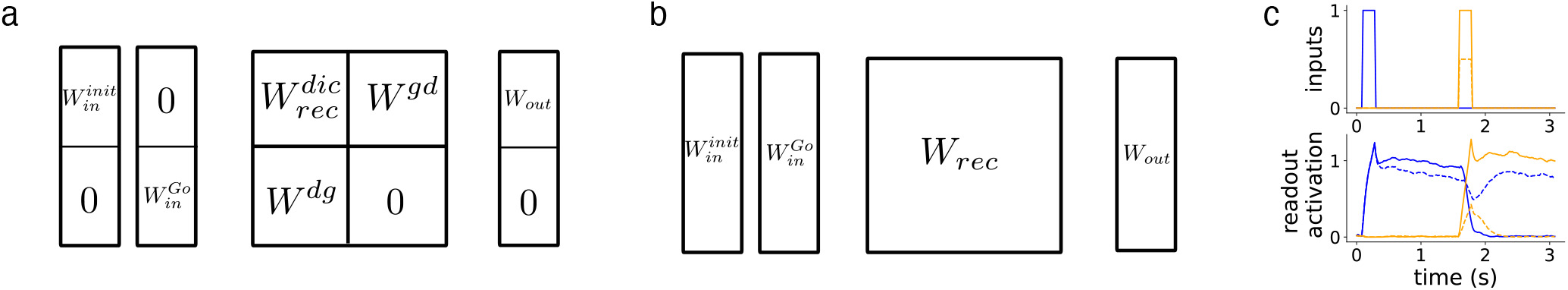
Training schemes for NLSC or for unconstrained ANN. **a**, Schematic of constraints on the connectivity for training NLSC, or **b** unconstrained networks. **c**, Top: inputs for two trials (plain and dashed lines) of the dictionary task. Bottom: activations of the blue and orange readout neurons for a trained networks in response to these inputs.

**SI Fig. 2:**
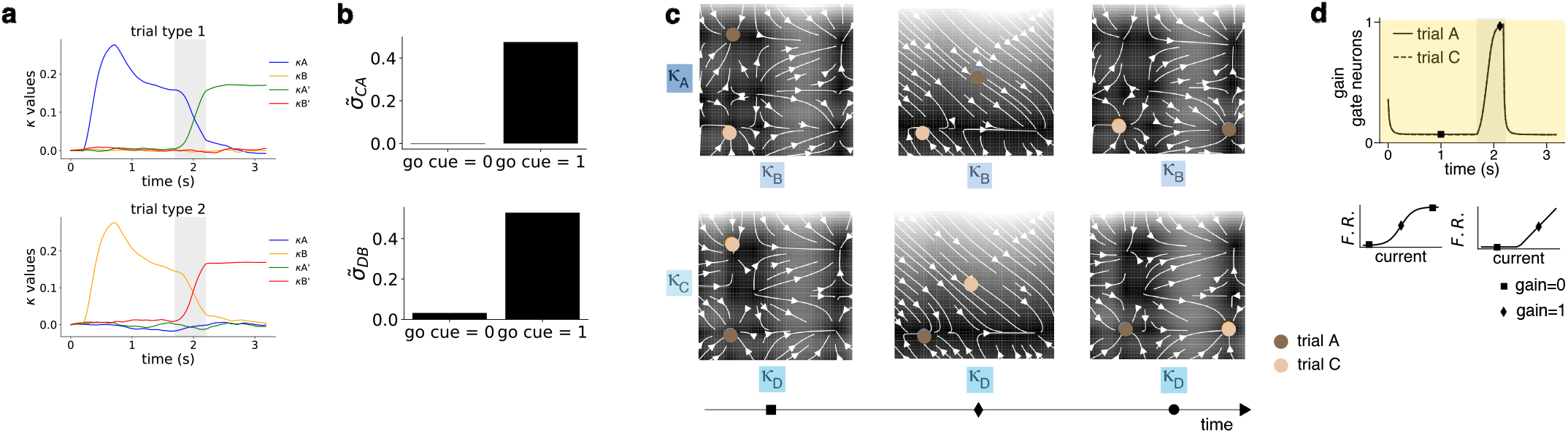
Reduced dynamical systems for NLSC. **a**, Sensory-motor transformation task: time evolution of the macroscopic variables describing the activation of cell assemblies during task performance. **b**, Interactions between macroscopic variables are modified by the go cue: activation transfers neural activity from state A to C or B to D. **c**, Flow fields for the 4-D reduced dynamical system of interacting cognitive variables. Light and dark brown dots represent current network state in trials A to B and C to D respectively. **d**, Flow-field is reshaped by setting the gain of a gate neurons from 0 to 1 in both types of trials.

**SI Fig. 3:**
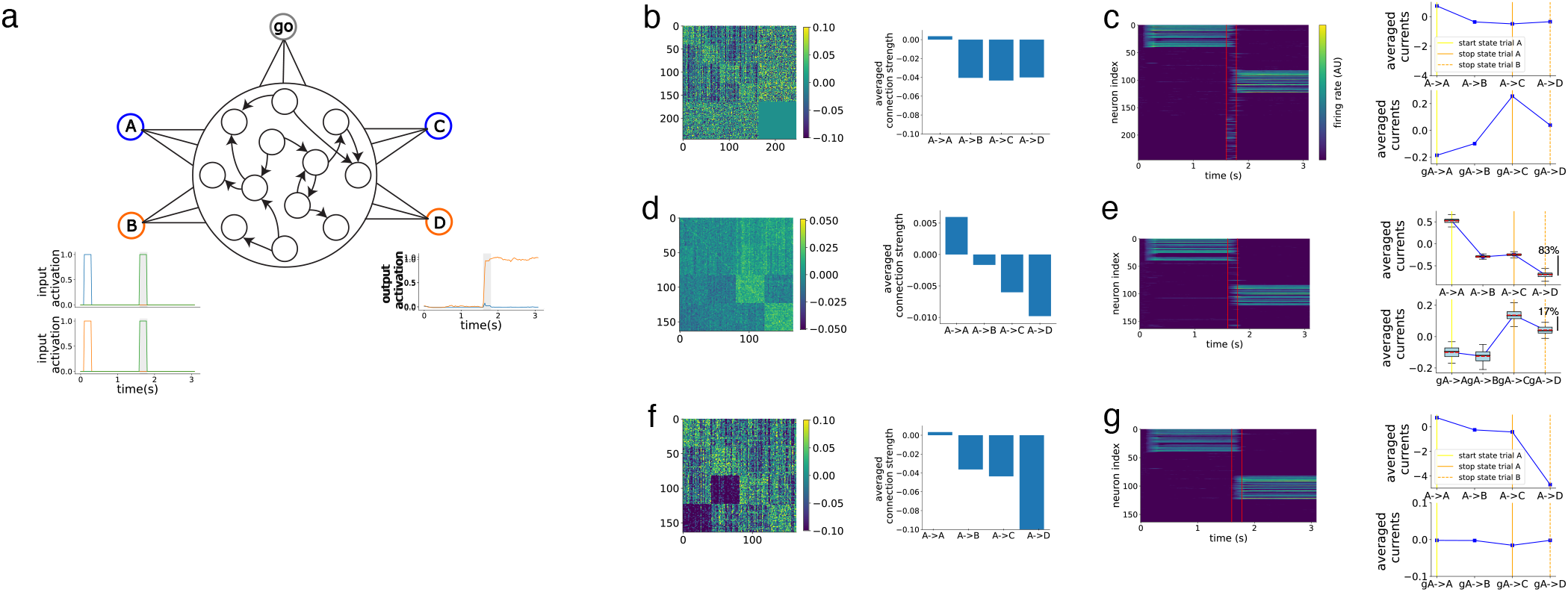
Neural Latching Switch Circuits in ANN trained on the delayed sensory-motor transformation task. **a**, Schematic of an ANN trained on the sensory-motor transformation task, below are inputs and outputs for the first (first row) and second (second row) trial type. **b**, Left: connectivity matrix of a NLSC trained on the sensory-motor transformation task. Right: average connection strengths over blocks of the matrix. First bar: connection strength among neurons belonging to the same cell assembly ; second bar: connection strength in between cell assemblies A and B, third bar: connection strength from cell assemblies A to C and B to D, fourth bar: connection strength from A to D and B to C. **c**, Left: raster of neural activity for a singe trial showing the activity of all neurons in the trained NLSC. Red lines materialize activation of the go cue. Right, top: averaged currents produced during the maintenance epoch by the maintenance state to: the maintenance state itself (e.g. A), the other potential maintenance state (e.g. B), the response state (e.g. C) and the other potential response state (e.g. D). Right, bottom: averaged currents produced during the go epoch by gate neurons. **d**, Same as **b** for unconstrained networks trained on the task. Left: example network, right: averaged over 50 networks. **e**, Same as **c**, with currents on the right averaged over the 50 networks. For the bottom plot, what we consider as gate neurons are the neurons that are not supposed to be active in that trial (e.g. neurons belonging to B and D for the first type of trials), but some of which are activated during the go epoch, see left. **f**,**g** Same as **b**,**c** for the network without gate neurons (see text).

**SI Fig. 4:**
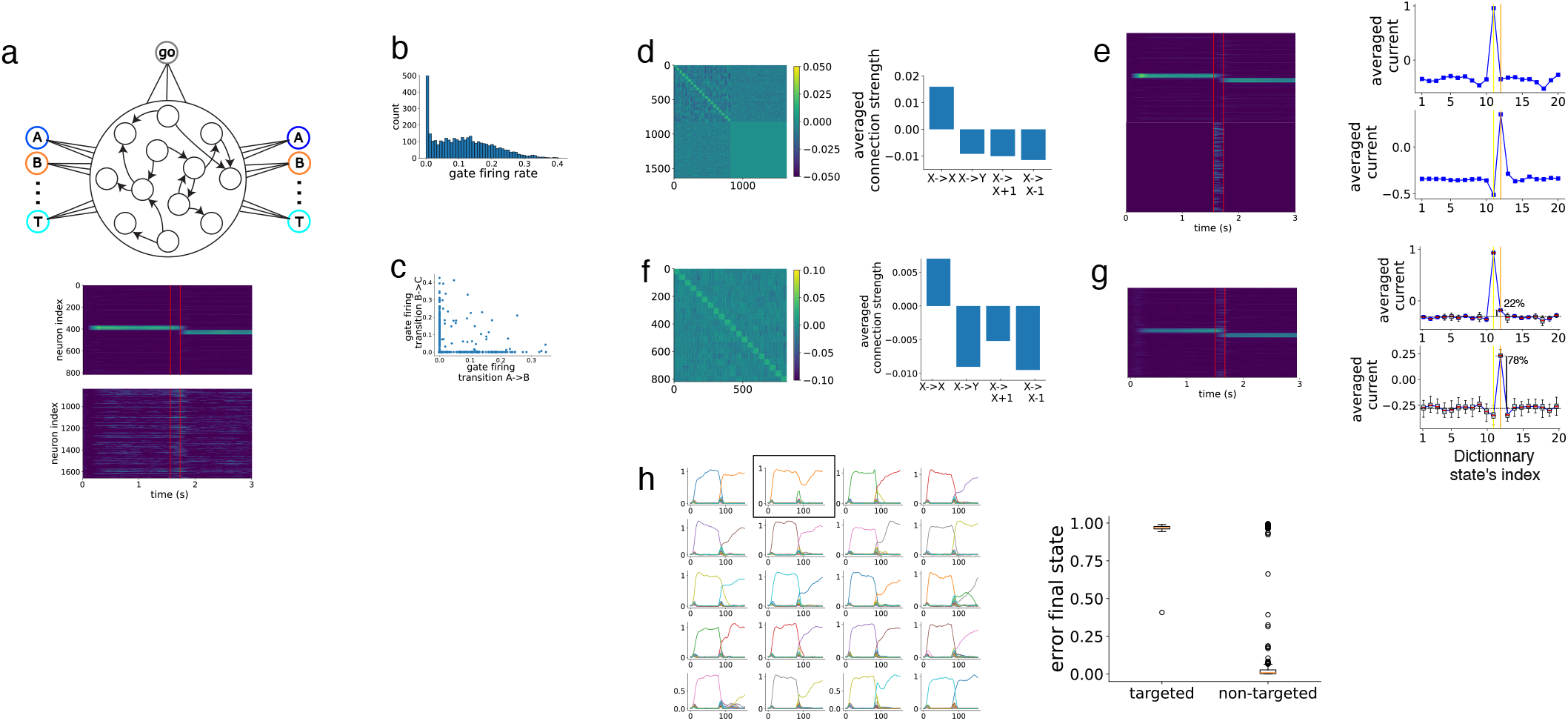
Neural Latching Switch Circuits in ANN trained on the permutation task. **a**, Schematic of an ANN trained on the permutation task, with raster of the NLSC with 820 dictionary and 820 gate neurons described in the main text. **b**, Distribution of activations for the 820 gate neurons for the 20 transitions, a gate neuron is considered active when its activity is above 0.05. **c**, Comparing activations of all gate neurons for two particular transitions. **d**, Left: connectivity matrix of a NLSC trained on the permutation task. Right: average connection strengths over blocks of the matrix. First bar: connection strength among neurons belonging to the same cell assembly ; second bar: connection strength in between two cell assemblies X and Y ≠ X, X-1, X+1 ; third bar: connection strength from X to X+1 ; fourth bar: connection strength from X to X-1. **e**, Left: raster of neural activity for a single trial showing the activity of all neurons in the trained NLSC. Red lines materialize activation of the go cue. Right, top: averaged currents produced during the maintenance epoch by the maintenance state to each of the states. The average is performed over neurons and over transitions. X-axis represents the index of the states shifted for each transition such that the maintenance state has index 11 (yellow line) and the response state has index 12 (orange line). Right, bottom: averaged currents produced during the go epoch by gate neurons to each of the states. **f**, Same as **d** for the averaged connectivity of 50 unconstrained networks trained on the permutation task. **g**, Same as **e** for the unconstrained networks. For the bottom plot, what we consider as gate neurons, for the transition from state X, are the neurons that belong to cell assembly Y ≠ X, X-1, X+1. **h**, Left: Readout activations for all the 20 transitions while inactivating 50 gate neurons during the go epoch. It prevents only the second transition (black square). The inactivated neurons are taken as the 50 with highest firing rate during the unperturbed transitions, with the constraint that they belong neither to the former, current, following activated cell assembly. Right: statistics of inactivations over all transitions. For each perturbed transition X, error in final state is measured as (z_X+1_(t = end) − 1)^2^.

**SI Fig. 5:**
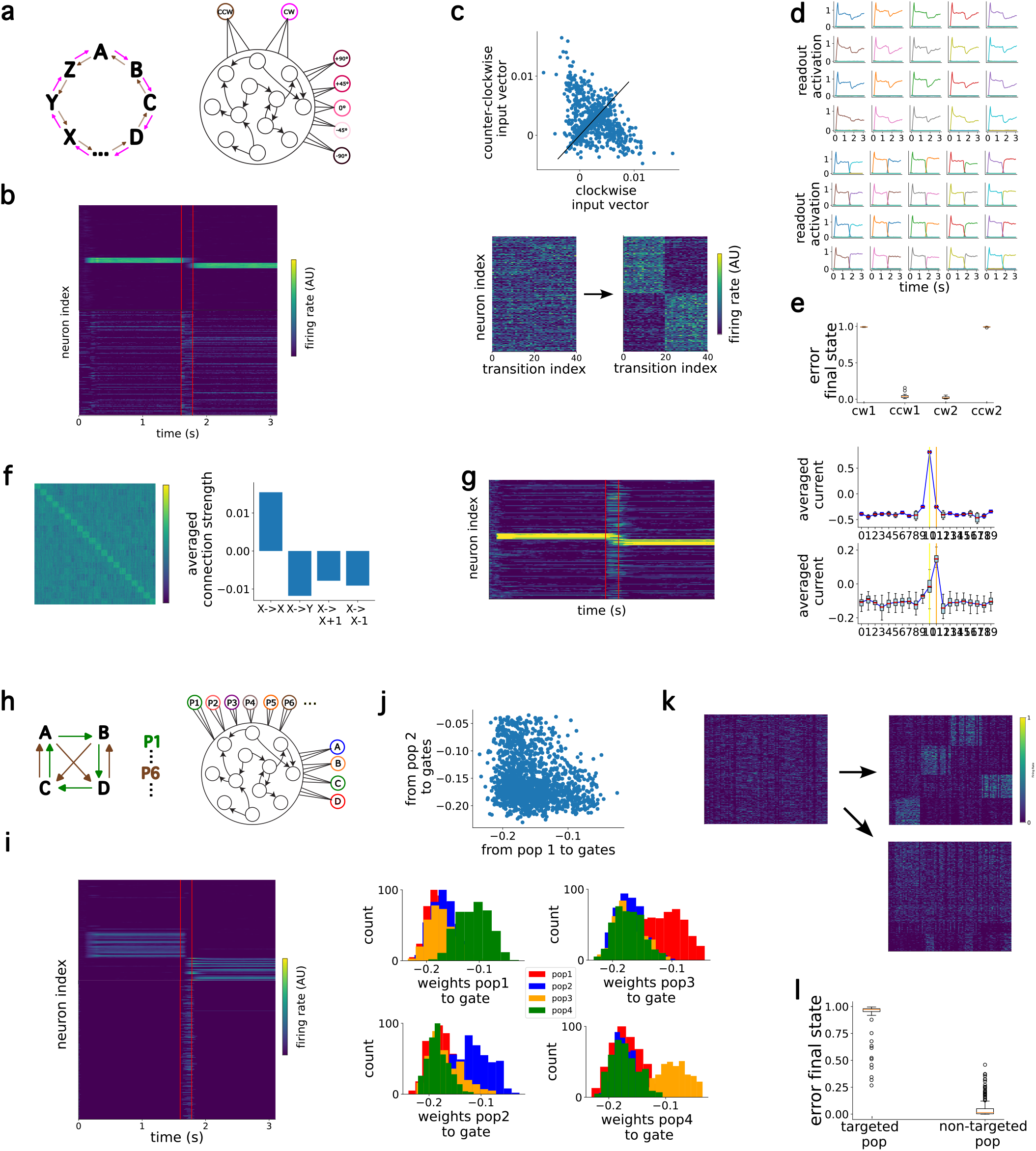
Head-direction system and multiple permutations over dictionary states. **a**, Schematic of an ANN trained on the head-direction task, where readout neurons and associated cell assemblies in the recurrent network encodes angular positions. **b**, For a single transition, raster of the NLSC with 820 dictionary and 820 gate neurons described in the main text. **c**, Top: individual entries of the two trained vestibular input connectivity vectors. Bottom: Reordering of the raster of activations of gate neurons for the clockwise and counter-clockwise transitions. **d**, Readouts activations for all the transitions (top clockwise, bottom counter-clockwise transitions) while inactivating the first population of gate neurons during the go epoch. **e**, Statistics of inactivation results for all transitions. The first two columns quantifies what is shown in (D), the next two represent inactivations of the second population of gate neurons. **f**, Averaged connectivity for 50 unconstrained networks trained on the head-direction task. **g**, Neural activity (left) and averaged currents generated by the maintenance state (right top) and transient activations during the go epoch (right bottom). **h**, Schematic of an ANN trained to implement all possible permutations of 4 states. **i**, For a single transition, raster of the NLSC with 164 dictionary and 1640 gate neurons described in the main text. **j**, Top: For individual gate neurons, averaged connectivity from dictionary neurons belonging to cell assembly encoding state 1 and state 2. Bottom: Each panel shows the histogram of averaged connectivity FROM dictionary neurons belonging to a cell assembly. Gate neurons are split into four groups (colors, see text). **k**, Reordering of the raster of gate neurons activations. Horizontal arrow shows reordering according to the clustering results in j, while the diagonal arrow shows reordering according to clustering gate neurons based on the averaged connectivity TO dictionary neurons belonging to a cell assembly. **l**, Statistics of inactivation results for all transitions and inactivation of one of the four populations. For a given transition, the inactivated population is either the one most activated (targeted pop), either one of the others (non-targeted pop).

**SI Fig. 6:**
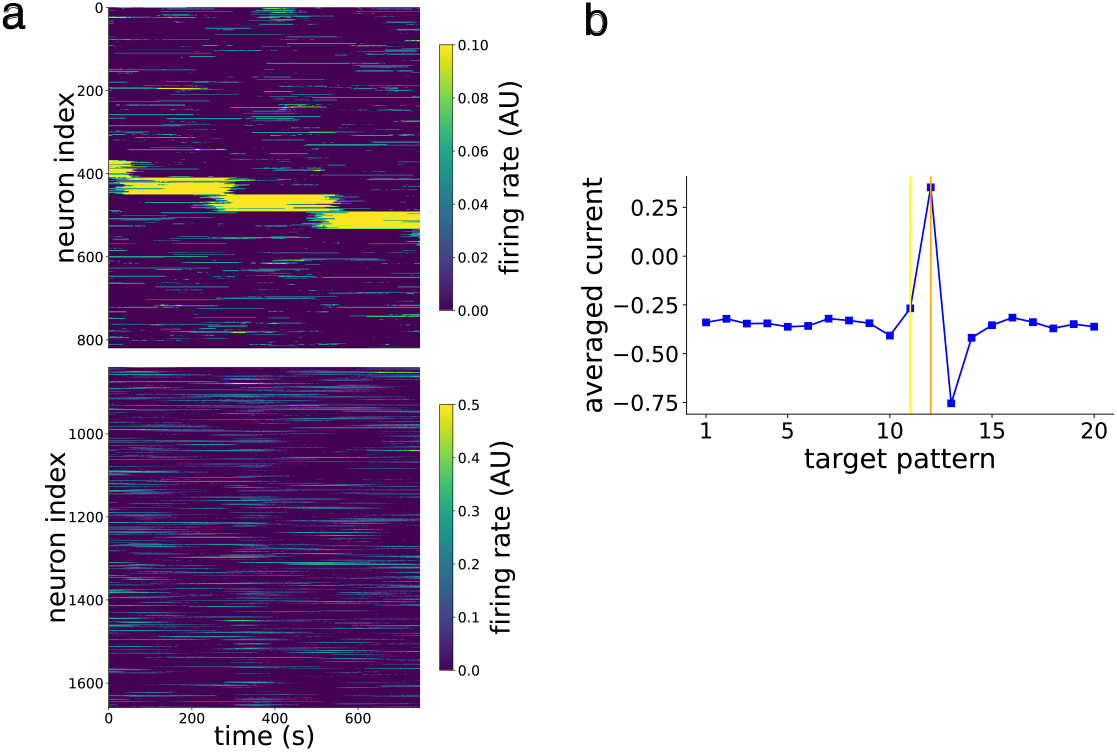
Relationship with oscillatory activity in a NLSC trained to autonomously update its dictionary states. **a**, Raster of activity for three transitions. **b**, Average current send from gate neurons to cell assemblies of the dictionary at transition time, averaged on all transitions, recentered on the transition from cell assembly 11 to cell assembly 12.

**SI Fig. 7:**
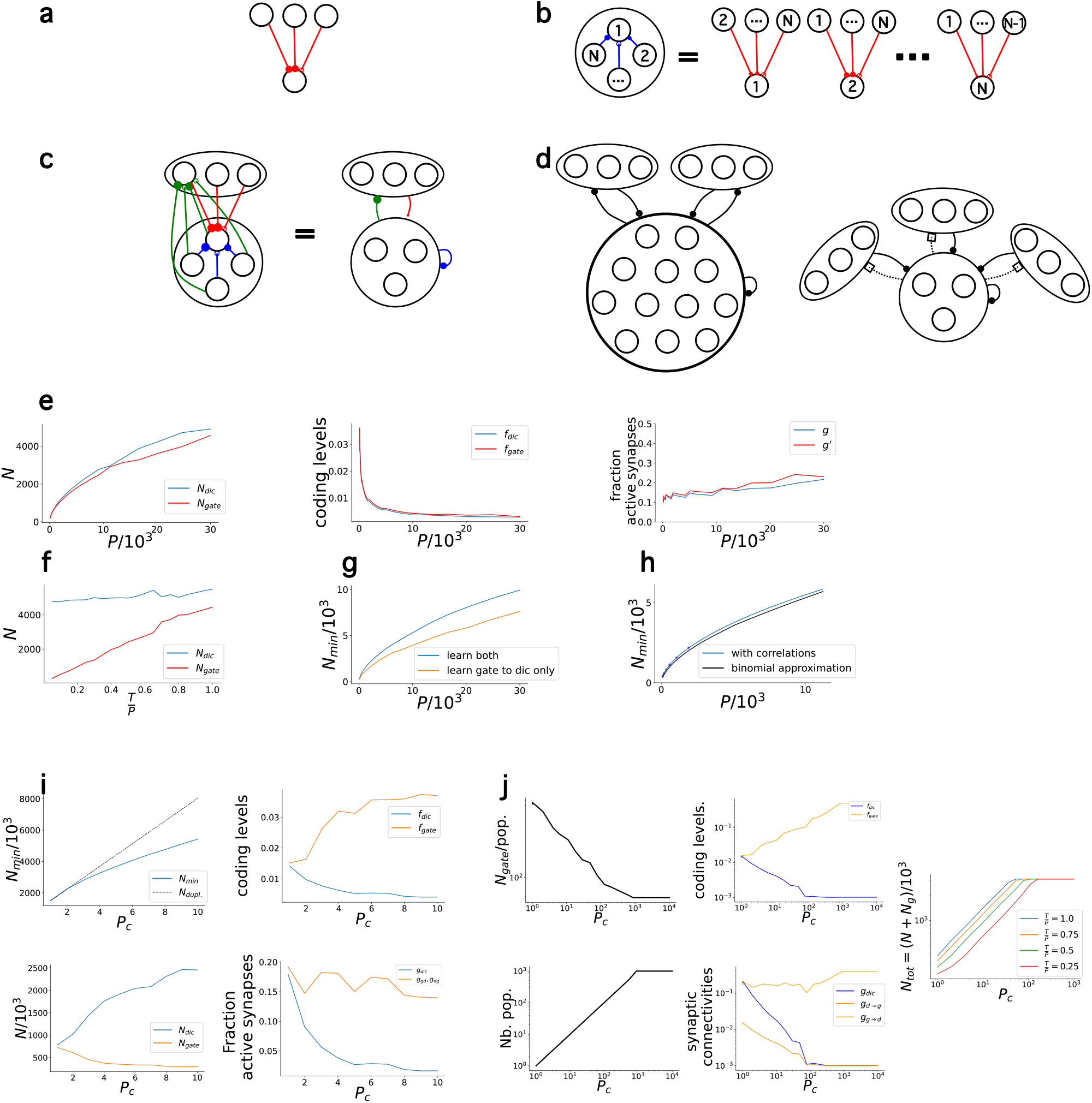
Capacity analysis of the NLSC. Cartoons for **a** a perceptron, **b** an attractor neural network, **c** a NLSC with a single population of gate neurons implementing a single permutation of dictionary states, **d** NLSC with multiple populations of gate neurons implementing multiple permutations of a large (left) or small (rigth) dictionary. **e**, Description of an optimized NLSC implementing a single permutation with an increasing number of dictionary states: number of dictionary and gate neurons (left), coding levels (middle), average synaptic weights (right). **f**, Number of dictionary and gate neurons as a function of the number of programmed transitions T. **g**, N_min_ for a network implementing a single permutation with learnt and random connectivity from dictionary to gate neurons. **h**, N_min_ as computed from **(8)** (blue) or **(10)** (black). **i**, Same as **e** for an optimized NLSC implementing an increasing number of permutations that remain small compared to dictionary size. **j**, Properties of optimized NLSC in the regimes of both low and large number of permutations per dictionary states.

## Footnotes

1 An alternative to this lowering of the activation threshold would be to have inhibitory connections from gate to dictionary neurons so that 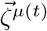 both activates 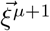 and silences 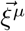.

